# *In Vivo* Dual RNA-Seq uncovers key toxin-like effectors of epithelial barrier disruption and tissue colonization by an extracellular bacterial pathogen

**DOI:** 10.1101/2025.04.04.647190

**Authors:** Giraud-Gatineau Alexandre, Haustant Georges, Monot Marc, Picardeau Mathieu, Benaroudj Nadia

**Affiliations:** Biology of Spirochetes, Institut Pasteur, Université Paris Cité, CNRS UMR 6047, F-75015 Paris, France; Plate-forme Technologique Biomics, Institut Pasteur, Université Paris Cité, F-75015 Paris, France

## Abstract

Disruption of host cell barriers is a fundamental strategy enabling pathogens to establish a paracellular infection. Using dual RNA-Seq, we determined the *in vivo* host-pathogen transcriptomic landscape upon infection by the extracellular pathogen *Leptospira interrogans* and uncovered a novel mechanism of cell-cell junction disruption. We demonstrated that, upon infection, an increase in intracellular calcium triggered tight junction destabilization, by activating the calmodulin and myosin light chain kinase signalization. We identified two novel bacterial effectors of the Virulence-Modifying (VM) proteins family, structurally related to toxin-like proteins, that promoted modulation of calcium homeostasis and disruption of cell-cell junctions, thereby allowing *Leptospira* translocation across epithelium barriers, tissue colonization and pathogenicity. Furthermore, we demonstrated that at least one of these VM proteins was internalized inside host cells. Altogether, these findings reveal a unique strategy by which an extracellular pathogen secretes toxin-like proteins to exploit host calcium signaling for breaching epithelial barriers.

## Main

Successful microbial infection and host colonization depend on the ability of pathogens to evade, resist and/or manipulate host defenses. While much of our understanding of mechanisms underlying the dynamic interplay between a host and a pathogen (i. e. host-pathogen interactions) stems from studies on intracellular pathogens, the mechanisms employed by extracellular pathogens remain comparatively underexplored. Moreover, these interactions have been generally investigated under *in vitro* condition that cannot fully replicate the complexity of a living host.

A critical feature to establish infection is the ability of extracellular pathogens to compromise the integrity of endothelium and epithelium barriers^1^, which are vital for maintaining organ integrity. Tissue integrity depends on protein scaffolds stabilizing cell-cell junctions, including tight junctions (TJs) and adherens junctions (AJs). TJs are formed by claudin, occludin and JAM-1 and zonula occludens (ZO) proteins while AJs are constituted by the calcium-binding protein cadherin^2^. Both TJs and AJs interact with the actin cytoskeleton to maintain barrier integrity. Pathogens often disrupt these junctions, directly or indirectly, through cytoskeletal regulation to invade a host^3^. The molecular mechanisms leading to cell-cell junction disruption and the bacterial effectors responsible for triggering this disruption remain poorly characterized upon infection by extracellular pathogens.

Among extracellular pathogens, *Leptospira* spp., the causative agents of leptospirosis, have evolved strategies to breach these barriers and disseminate throughout the host. These bacteria enter through skin abrasion or mucosa, evade the innate immune response and rapidly disseminate hematogenously to organs, including kidneys and liver, resulting in multiple organ failure and hemorrhage. With an estimated one million severe cases and 60,000 deaths reported annually^4^, leptospirosis is the most widespread zoonotic disease. Previous studies have demonstrated that *L. interrogans* infect their mammalian host through a paracellular route, disrupting both TJs and AJs^5,6^. Because *Leptospira* exclusively employs a paracellular route to colonize host tissues, it served as an ideal model to understand mechanisms of cell-cell junction disruption by extracellular pathogens.

Here, we developed a dedicated protocol for dual RNA-Seq profiling of *L. interrogans*-infected hamsters, a model of human acute leptospirosis, to elucidate how the extracellular pathogen *Leptospira* spreads and compromises cell barrier integrity. Transcriptional host response was characterized by alterations in pathways associated with paracellular permeability, including cell-cell junctions and cytoskeletal organization. We demonstrated that the disruption of cell-cell junctions by *L. interrogans* correlated with calcium homeostasis deregulation and activation of calmodulin and MLCK kinases. Obtaining the *in vivo L. interrogans* transcriptomic profile allowed for the identification of two leptospiral toxin-like proteins of the Virulence-Modifying (VM) family. We established their essentiality in *Leptospira* pathogenicity and their involvement in regulating the calcium-induced calmodulin/MLCK signaling pathway, leading to cell-cell junction disassembly and promoting paracellular transmigration through host tissues. These findings revealed a unique mechanism by which an extracellular pathogen alters host calcium signaling to disrupt cell barriers for facilitating its dissemination and colonization inside the host.

## Results

### Dual RNA-Seq of *L. interrogans*-infected hamsters

To define the optimal condition for performing dual RNA-Seq in the model of human acute leptospirosis, Golden Syrian hamsters were infected with 10^8^ *L. interrogans* by intraperitoneal route. In these conditions, colonization of *L. interrogans* was detectable in liver and kidneys at 1-day post-infection (dpi), with signs of morbidity apparent at 4 dpi (**Fig. 1A-B**). To assess transcriptional changes at early and late stages of infection, we performed RNA sequencing of the host and *Leptospira* on liver and kidney tissues collected at 1 and 3 dpi (**Fig. 1C**). Initial sequencing, performed at standard read depths (∼40 million reads/sample), revealed that more than 99.9% of total reads mapped to hamster genome, allowing for the identification of up to 984 differentially expressed genes (DEGs, *p.adj* < 0.05; |Log2FC| >1) (Fig 1D, **Extended data Fig. 1A**). Among all conditions tested, the highest proportion of *Leptospira* reads was obtained by sequencing RNAs isolated from liver tissue at 3 dpi, detecting the expression of only 85 *Leptospira* genes (**Extended data Fig. 1A**). To overcome this limitation and achieve higher *Leptospira* transcriptome coverage, RNAs isolated from liver at 3 dpi were deep-sequenced (∼400 million reads/sample). This approach significantly increased the proportion of *Leptospira* genes detected from 4,57% to 79.34%, with a robust and well-distributed read count across *Leptospira* genome (**Fig. 1E, Extended data Fig. 1B**). Thus, this experimental design provides a comprehensive and robust dual RNA sequencing profile encompassing both the host and pathogen transcriptomic responses.

**Figure 1.**
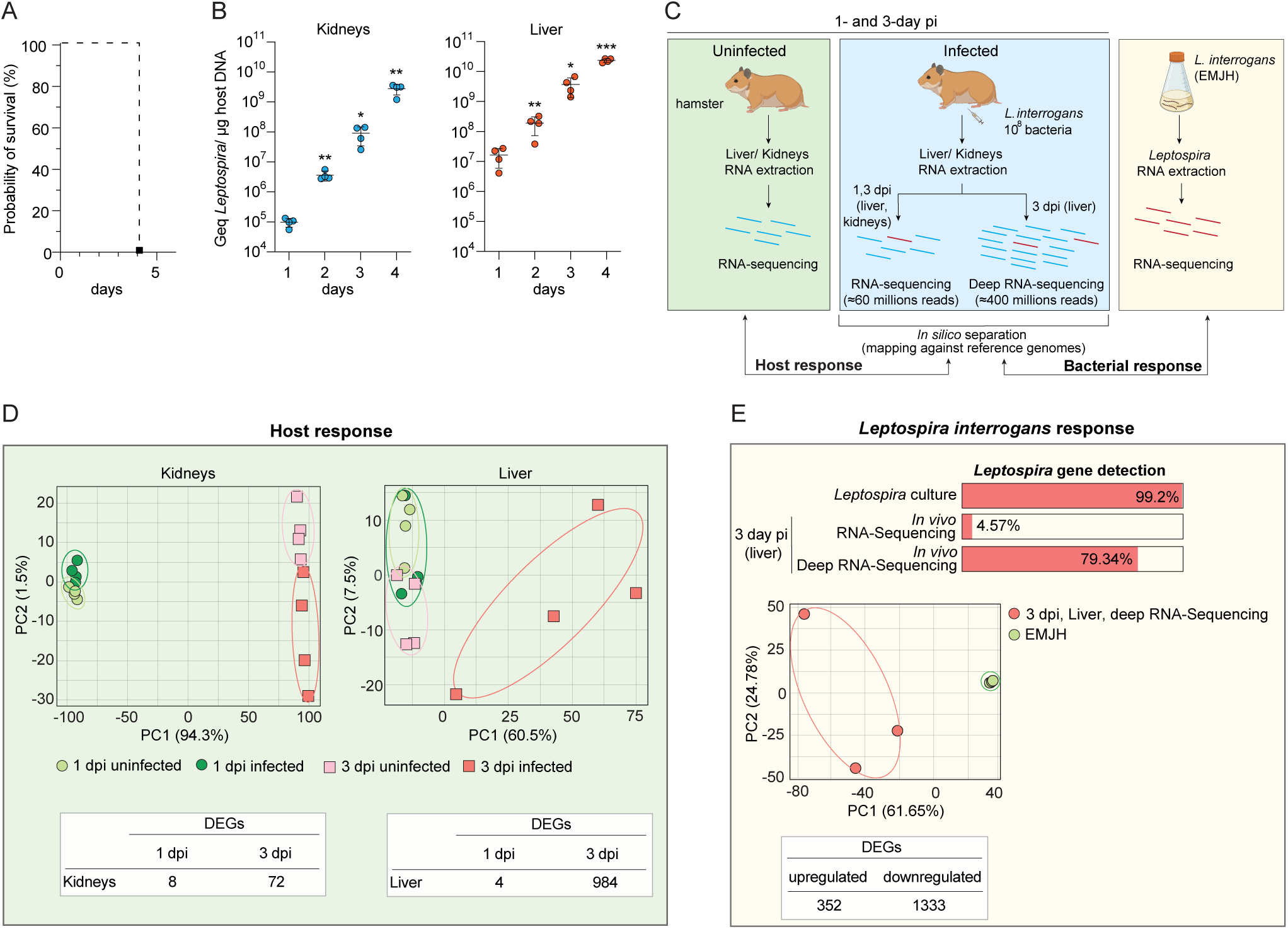
Design and evaluation of the *in vivo* Dual RNA-seq for *L. interrogans*-infected hamsters (A) Survival curve of hamsters infected with *L. interrogans*. The virulence *of L. interrogans* serovar Manilae was evaluated in a group of four hamsters via an intraperitoneal infection with 10^8^ leptospires. (B) Leptospiral burden in liver and kidneys was quantified by qPCR analysis using the *L. interrogans flaB2* and hamster *gadph* genes at 1-, 2-, 3-and 4-day post-infection and expressed at genomic equivalents (GEq) per μg of hamster DNA. Data are the mean ± SD and statistical significance was determined by unpaired two-tailed Student’s *t* test compared to 1 dpi. *, *p* < 0.02; **, *p* <0.002; ***, *p* <0.0002. (C) Schematic description of the dual RNA-seq workflow. Hamsters (n= 4) were infected with 10^8^ *L. interrogans* by the intraperitoneal route. At 1 and 3 dpi, RNAs were extracted from liver and kidney tissues, and sequenced. In parallel, RNA extraction and sequencing were performed on exponentially growing *L. interrogans* cultivated in EMJH medium. Host responses were analyzed by comparing transcriptomes of infected versus uninfected tissues. A deep RNA-sequencing was performed on samples isolated from liver at 3 dpi (n= 3) to enrich bacterial reads and enable comparison of *L. interrogans* transcriptional profiles *in vivo* and *in vitro*. Hamster and leptospiral RNAs were represented in blue and red, respectively. (D) Principal component analysis (PCA) of the global gene expression in liver (right panel) and kidney (left panel) tissues upon *L. interrogans* infection at 1 (green circles) and 3 (red squares) dpi. Uninfected conditions (used as controls) are presented by light green (1 dpi) and light red (3 dpi) symbols. The numbers of differentially expressed genes (DEGs) (FDR ≤0.05; |Log2FC| ≥1) in liver and kidneys at 1-and 3-dpi are indicated for each condition. (E) Proportion (expressed as percentage) of *L. interrogans* genes detected in each dataset (at 3 dpi), highlighting the enrichment achieved through deep sequencing, and PCA of the global gene expression of *L. interrogans* in liver at 3 dpi (red symbols) compared to EMJH condition (green symbols). The numbers of leptospiral DEGs (FDR ≤0.05; |Log2FC| ≥1) in hamster liver compared to EMJH condition are indicated.

### Transcriptomic host response upon *Leptospira* infection

Principal component analysis (PCA) of the RNA-Seq data set revealed distinct clustering of infected compared to uninfected samples at 3 dpi, while no separation was observed at 1 dpi, in liver as well as in kidney tissues (**Fig. 1D**). Despite the detection of a significant *L. interrogans* load at 1 dpi (**Fig. 1B**), the host transcriptional response was minimal, with only 4 and 8 DEGs (*p.adj* < 0.05; |Log2FC| >1) in the liver and kidneys, respectively (**Fig. 1D, Source data Fig. 2**). By 3 dpi, the transcriptional response was increased significantly, with 984 and 72 DEGs (*p.adj* < 0.05; |Log2FC| >1) in the liver and kidneys, respectively (**Fig. 1D, Extended data Fig. 2A, Source data Fig. 2**). These findings indicate that *L. interrogans* do not elicit an early host transcriptional response, even at a high infectious dose (10^8^), given a LD50 of 10-100 leptospires^7^.

**Figure 2.**
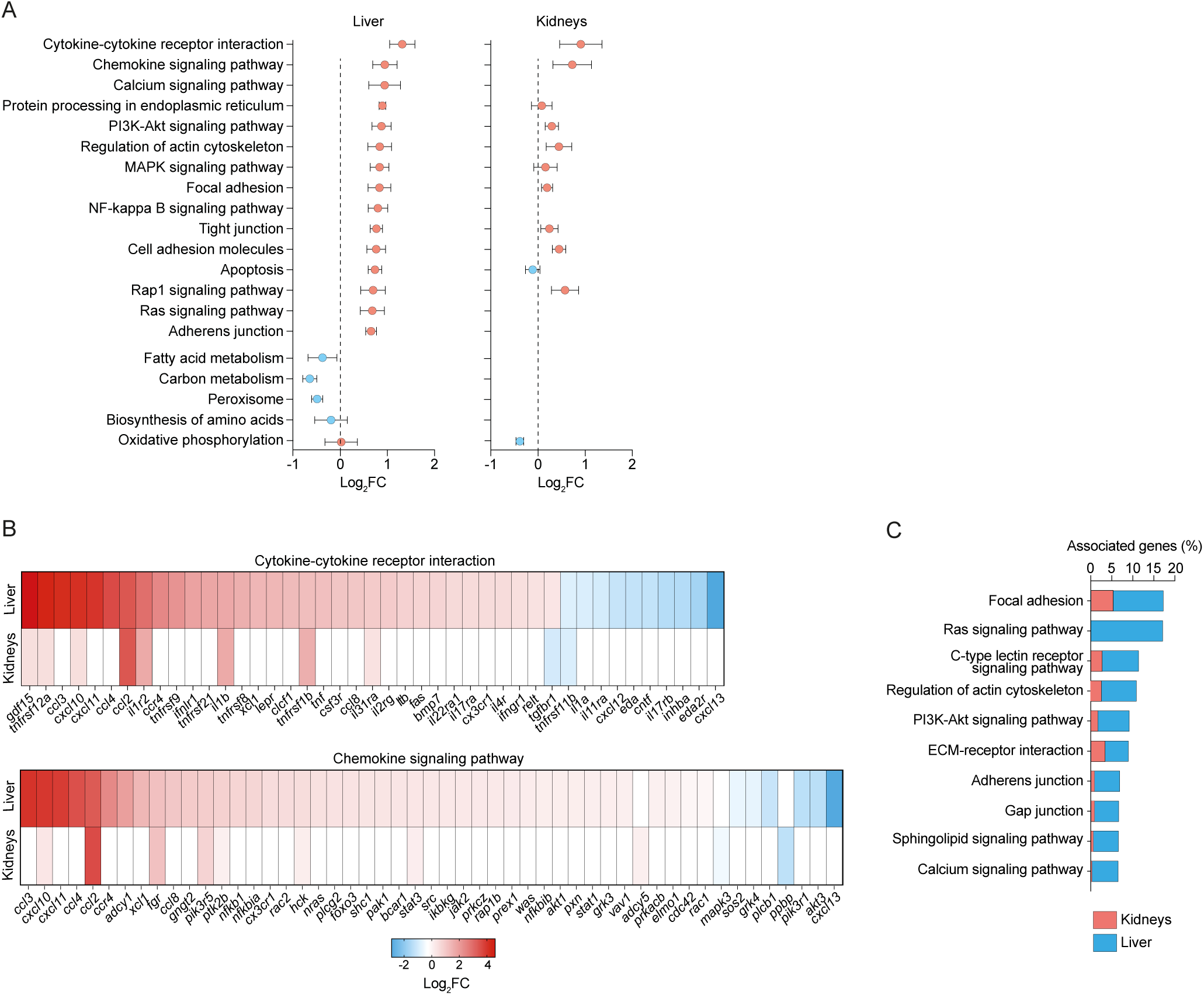
Host transcriptional response to *L. interrogans* infection *in vivo*. (A) KEGG analysis of major upregulated and downregulated pathways in liver and kidneys at 3 dpi. The y-axis shows the enriched pathways and the x-axis values are the mean Log2FC of the DEGs in each enriched pathway. Up-and downregulated ORFs are represented by red and blue symbols, respectively. (B) Heatmap showing DEGs related to Cytokine-cytokine receptor interaction (upper panel) and Chemokine signaling pathway (lower panel) in liver and kidneys upon *L. interrogans* infection. Gene names are indicated at the bottom of the panels. Differential expressions are expressed as Log2FC (infected versus uninfected) with a heat map color from blue to red indicating low to high Log2FC. (C) KEGG pathways analysis of DEGs in *L. interrogans*-infected hamsters involved in cytoskeleton organization Gene Ontology group. Only the ten most significative pathways are represented (with a *p*-value corrected with Bonferroni step down < 0.005). The percentage of DEGs associated with each pathway is indicated for the liver (red) and kidneys (blue).

At 3 dpi, both liver and kidney exhibited transcriptional changes associated with inflammation, cell-cell junctions, and cytoskeletal organization (**Fig. 2A, Extended data Fig. 2 and Extended data Table 1**). Pathway enrichment analysis revealed upregulation of cytokine-cytokine receptor interaction (e.g., *gdf15*, *ccl2*, *cxcl10*, *il1b*) and chemokine signaling (*fgr*, *hck*, *stat3*), highlighting a common inflammatory response in both organs (**Fig. 2B**). Genes involved in the actin cytoskeleton, focal adhesions, tight junctions, and cell adhesion were also upregulated, including genes encoding integrins (*itgal* in both liver and kidney, *itgb6*, *itgav*, *itga5*, *itga3* in liver, and *itga4*, *itga8* in kidney), as well as actin-related proteins (*mlck*, *actg1* in liver; *mapk3*, *pip5k1n* in kidneys; **Fig. 2C, Extended data Fig. 2 and Extended data Table 1**)

In addition to the transcriptional responses shared by the liver and kidneys, organ-specific differences emerged. In the liver, transcriptional changes were more pronounced, with strong upregulation of inflammatory pathways, protein processing in the endoplasmic reticulum and cell-cell junction pathways (**Fig. 2A, Extended data Fig. 2 and Extended data Table 1**). In addition, we observed an upregulation of the calcium signaling pathway and genes encoding factors related to cell-cell junction integrity such as cadherin (*cdh1*) and tight junction protein (*tjp2*, encoding ZO-2). In contrast, the kidneys showed a more restricted transcriptional response, with fewer differentially expressed genes (**Fig. 1D**). A key distinction was the downregulation of specific metabolism-related pathways in the two organs. The liver response showed a specific downregulation of amino acid and fatty acid metabolism, whereas the kidney response was associated with a downregulation of oxidative phosphorylation (**Fig. 2A, Extended data Fig. 2 and Extended data Table 1**). Taken together, these findings indicate that, upon leptospiral infection, while both organs activate the inflammatory response and cell-cell junction pathways, the liver undergoes a more extensive transcriptional shift.

### *Leptospira* transcriptional profile *in vivo*

Analysis of the transcriptomic response of *L. interrogans* in liver of infected hamsters showed that the expression of 1685 *Leptospira* genes (59% of total ORFs) was affected *in vivo* compared to *in vitro* condition, with 352 and 1333 up-and downregulated genes (*p.adj* <0.05; |Log2FC| >1), respectively (**Fig. 3A, Source data Fig. 3**). Pathway enrichment analysis indicated that upregulated genes were predominantly linked to bacterial secretion system, O-antigen biosynthesis and key metabolic pathways such as carbon, amino acid, and porphyrin metabolisms (**Fig. 3B**, Extended Data Fig. 3 a**nd Extended data Table 2**). Notably, several genes encoding molecular chaperones involved in the proteotoxic stress response, such as *groES, groEL*, *hsp15*, *hsp20*, *dnaK*, *dnaJ*, and *grpE* were among the most highly upregulated (**Fig. 3C**). Additionally, the expression of several virulence-associated genes was significantly increased, including two genes encoding members of the Virulence-Modifying (VM) family, *sph* genes (encoding sphingomyelinase C), *ligA* and *ligB* (coding for immunoglobulin-like proteins A and B), *colA* (encoding collagenase A) and *LIMLP_09380* (encoding a hypothetical membrane protein)^8^.

**Figure 3.**
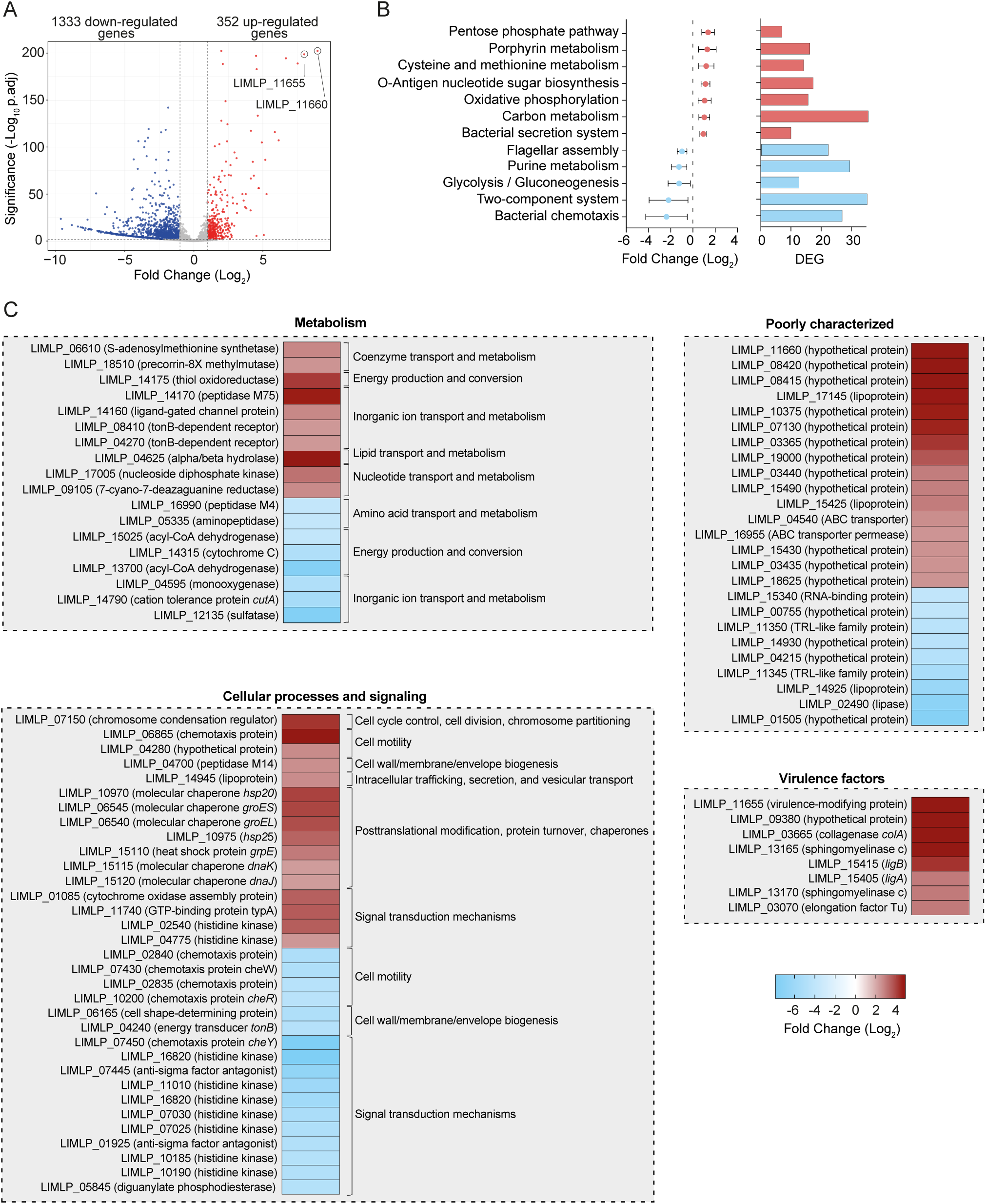
Transcriptional landscape of *L. interrogans in vivo* (A) Volcano representation of *L. interrogans* DEGs in liver of infected hamsters (n=3) at 3 dpi compared to *L. interrogans* cultivated in EMJH medium (in triplicate). Up-and downregulated ORFs are represented in red and blue, respectively, genes that are not significantly differentially expressed are shown in grey (FDR ≤0.05; |Log2FC| ≥1). The locus name of the two most upregulated genes under *in vivo* condition is indicated. (B) KEGG analysis of the most deregulated pathways in *L. interrogans* in liver of infected hamsters at 3 dpi. The *y*-axis shows enriched pathways and values of the *x-*axis are the mean of Log2FC of DEGs for each enriched pathway. The number of DEGs for each pathway is represented in the right panel. Up-and downregulated ORFs are represented in red and blue, respectively. (C) Heatmap of selected up-and downregulated genes under *in vivo* condition (*L. interrogans* DEGs in liver of infected hamsters at 3 dpi versus *L. interrogans* cultivated in EMJH medium). Genes were classified using COG annotation. Gene names and nomenclatures are indicated on the left of each panel. Differential expressions are expressed as Log2FC (*L. interrogans* in liver of infected hamsters at 3 dpi versus *L. interrogans* cultivated in EMJH medium) with a heat map color from blue to red indicating low to high Log2FC.

By contrast, genes involved in flagellar assembly, bacterial chemotaxis and two-component systems were downregulated (**Fig. 3B**, **Extended Data Fig. 3**). This included the decreased expression of a cluster (*LIMLP_07420-07460*) containing several genes associated with chemotaxis, such as *mcpa* and *cheABDRWY* and of several histidine kinase-encoding genes (*LIMLP_11010*, *LIMLP_10185*, *LIMLP_10190*, *LIMLP_16820*) (**Fig. 3C**). In addition, we observed the downregulation of genes associated with lipid, carbohydrate and coenzyme transport and metabolism (**Extended Data Table 2**).

Altogether, these findings revealed a major transcriptional reprogramming of *L. interrogans in vivo*, highlighting stress adaptation, metabolic shift and virulence induction as key features of adaptation to the host environment.

### *L. interrogans* induces actin reorganization and disrupts epithelial cell junctions in the host

The host transcriptional response suggested cell-cell junction regulation during leptospiral infection. We evaluated whether changes in expression of genes involved in this process correlated with F-actin rearrangement and cell junction disruption in human epithelial cells infected by *L. interrogans*. Live-cell and immunofluorescence microscopy showed that human epithelial cells infected with *L. interrogans* exhibited dispersion and a significant disorganization of actin filaments at 24 hr pi, resulting in reduced cell-cell contacts (**Fig. 4A-C**). Such phenomenon was not observed in cells infected with the saprophytic species *L. biflexa* (**Extended Data Fig. 4**). Importantly, no epithelial cell death was observed for at least 48 hr pi (**Fig. 4D**), indicating that actin rearrangement was not related to cell death. Immunoblot analysis showed increased cellular contents of actin nucleation and polymerization factors (Profilin-1, WAVE-2, and phosphorylated-Rac1), as well as of the actin reorganization factor phosphorylated Cofilin (**Fig. 4E**). Of note, genes encoding these factors were upregulated upon *Leptospira* infection in liver (**Extended Data Table 3**).

**Figure 4.**
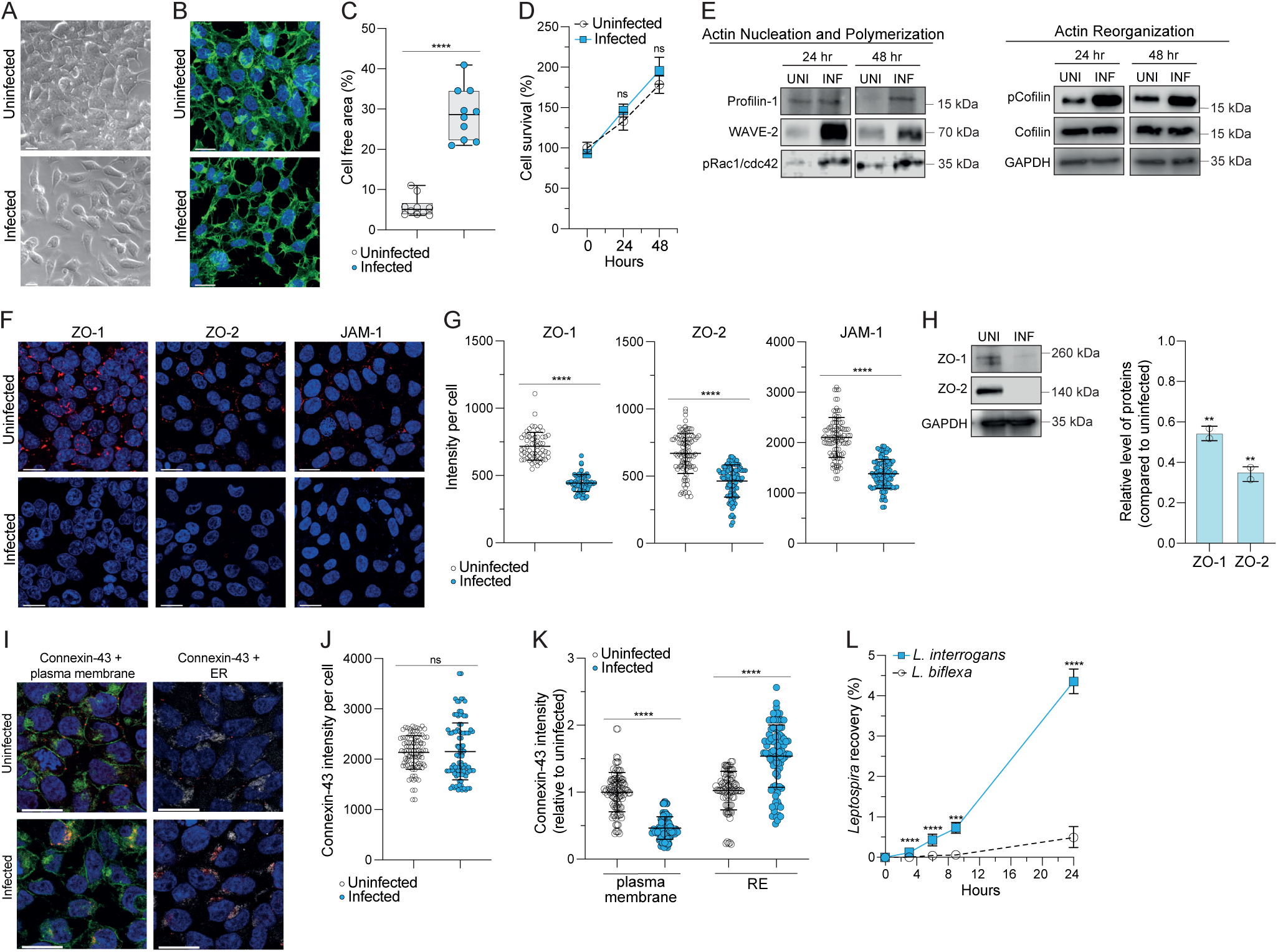
*L. interrogans* induces actin remodeling and disrupts cell-cell junctions in human epithelial cells. (A) Phase contrast images of human epithelial cells (HEK293T) uninfected (upper panels) or infected with *L. interrogans* for 24 hr (lower panels). Scale bars, 20 µm. (B) Confocal images of human epithelial cells uninfected (upper panels) or infected with *L. interrogans* for 24 hr (lower panels) and stained for F-actin (in green) and DNA (in blue). Scale bars, 10 µm. The images are representative of 3 biological replicates. (C) Quantification of the percentage of cell-free area for uninfected (empty symbols) or infected condition (blue symbols) (n=10 fields of view). Box and whisker plots represent the median and the interquartile range ± min and max values, respectively, from 3 independent biological experiments. ****, *p* < 0.0001 (unpaired, tailed *t* test). (D) Cell viability was assessed at 0, 24 and 48 hr in uninfected (empty circles) and *L. interrogans*-infected (blue squares) cells, from 3 independent biological experiments (unpaired, tailed *t* test). (E) Immunoblot detection of Profilin-1, WAVE-2, pRac1/cdc42, pCofilin, and Cofilin in total lysates from HEK293T cells uninfected (UNI) or infected (INF) for 24 and 48 hr with *L. interrogans*. Immunoblots were performed as described in the Methods section. GAPDH cellular content was used as an equal loading control. The molecular mass markers (in kilodaltons) are indicated on the right. Data are representative of 2 independent experiments. (F) Confocal images of human epithelial cells uninfected (upper panels) and infected with *L. interrogans* for 24 hr (lower panels), stained for ZO-1, ZO-2 or JAM-1 (in red) and DNA (in blue). Scale bars, 20 µm. The images are representative of 3 biological replicates. (G) Fluorescence quantification of ZO-1, ZO-2 and JAM-1 intensity per cell (n = 100 cells) in infected (blue symbols) or uninfected (empty symbols) samples. ****, *p* < 0.0001 (unpaired, tailed *t* test). (H) ZO-1, ZO-2 and JAM-1 cellular content of uninfected (UNI) and infected (INF) cells as assessed by immunoblot. GAPDH cellular content was used as an equal loading control. The molecular mass markers (in kilodaltons) are indicated on the right. The quantified relative protein contents in infected cells (normalized by the values in uninfected controls) are plotted on the right panel. Data are mean ± SD from 2 independent experiments. **, *p* < 0.05 (unpaired, tailed *t* test). (I) Confocal images of human epithelial cells uninfected (upper panels) and infected with *L. interrogans* for 24 hr (lower panels) stained for Connexin-43 (in red), plasma membrane (in green, left panels), ER (in white, right panels) and DNA (in blue). Scale bars, 20 µm. The images are representative of 3 biological replicates. (J) Fluorescence quantification of Connexin-43 intensity per cell (n = 100) in epithelial cells uninfected (empty symbols) and infected (blue symbols); (unpaired, tailed *t* test). (K) Ratio of Connexin-43 colocalizing with plasma membrane or ER (as indicated). Values obtained for infected cells (blue symbols) were normalized by those in uninfected cells (empty symbols) (n = 100 cells). ****, *p* < 0.0001 (unpaired, tailed *t* test). (L) Ability of *L. interrogans* (blue squares) and *biflexa* (empty circles) strains to translocate through human epithelial cells. Bacteria were inoculated in the upper chamber of a cell culture insert containing human epithelial cell monolayer. *L. biflexa* were used as negative controls. The percentage of *Leptospira* recovery in the lower chamber was determined by qPCR analysis. Values obtained at the indicated times were normalized by that of the inoculum and expressed as percentage. Data are mean ± SD from 3 independent experiments. ***, *p* < 0.001; ****, *p* < 0.0001 (unpaired, two-tailed *t* test).

In addition, infected cells exhibited at 24 hr pi a reduction in tight junction proteins ZO-1, ZO-2, and JAM-1 compared to uninfected controls (1.5-, 1.4-and 1.6-fold reductions for ZO-1, ZO-2 and JAM-1, respectively, as measured by immunofluorescence, **Fig. 4F-G**). Such decreases were also observed by immunoblot (**Fig. 4H**). Interestingly, infection by *L. interrogans* did not lead to a decrease of the gap junction protein Connexin-43 cellular content but to a change in its localization. Indeed, while Connexin-43 was primarily localized at the plasma membrane in uninfected cells, it accumulated in the endoplasmic reticulum (ER) in infected cells (**Fig. 4I-K**). Finally, disruption of cell junctions upon infection by *L. interrogans* correlated with a greater ability to cross epithelial monolayers, compared with the saprophytic *L. biflexa* which lacks the capacity to disrupt cell-cell junction integrity (**Fig. 4L**). Altogether, these results demonstrate that infection by *L. interrogans* leads to a profound reorganization of the actin cytoskeleton and compromises epithelial barrier integrity probably by triggering downregulation or mislocalization of tight and gap junction proteins.

### ER stress response and its implications in *Leptospira*-infected cells

The host transcriptional response revealed an upregulation of genes associated with the ER stress pathway during infection by *L. interrogans* (**Fig. 2A** and **Supplementary Fig. 1A**). Upregulation of selected ER-related genes was confirmed *in vitro* by RT-qPCR (**Supplementary Fig. 1B**). Given the correlation between ER stress and the disruption of cell-cell junctions^9–11^, we evaluated whether change in expression of ER-related genes correlated with modulation of ER activity upon infection by *L. interrogans* using a fluorescent marker specific to the ATP-sensitive K^+^ channels of this organelle. At 6, 24 and 48 hr pi, the fluorescence intensity was significantly reduced in infected cells compared to uninfected control cells (**Supplementary Fig. 1C-D**), suggesting an alteration of ER function. Moreover, the phosphorylation of two ER stress markers, i. e. PERK and IRE-1, was increased in infected cells (**Supplementary Fig. 1E**), indicating activation of the ER stress response. To determine whether this ER stress response contributed to cell junction disruption, infected cells were treated with different ER stress inhibitors. However, none of the inhibitors restored cell-cell junction integrity in *L. interrogans*-infected cells (**Supplementary Fig. 1F-G**). Therefore, while *L. interrogans* induces an ER stress response probably mediated by PERK and IRE-1 pathways, this response does not substantially contribute to cell-cell junction disruption.

### Calcium signaling participates in cell-cell junction disruption in *L. interrogans*-infected cells

Given the upregulation of calcium-related genes upon *L. interrogans* infection (**Fig. 2C**) and the established role of Ca^2+^ signaling in cytoskeletal dynamics^12^, we investigated whether this cation contributed to cell-cell junction disruption. Intracellular Ca^2+^ levels increased by 1.65-and 1.39-fold in infected epithelial cells at 24 and 48 hr pi, respectively, compared to uninfected controls (**Fig. 5A-B**). To determine whether this Ca^2+^ accumulation was functionally linked to junction disruption, we treated infected cells with BAPTA-AM, a cell-permeable Ca^2+^ chelator. This treatment effectively limited the infection-induced Ca^2+^ increase, reduced cell junction dispersion and increased ZO-1 localization at cell-cell junctions compared to untreated infected cells (**Fig. 5C-E, Extended Data Fig. 5**). These results suggest that intracellular Ca^2+^ plays an important role in *Leptospira*-induced junction disruption.

**Figure 5.**
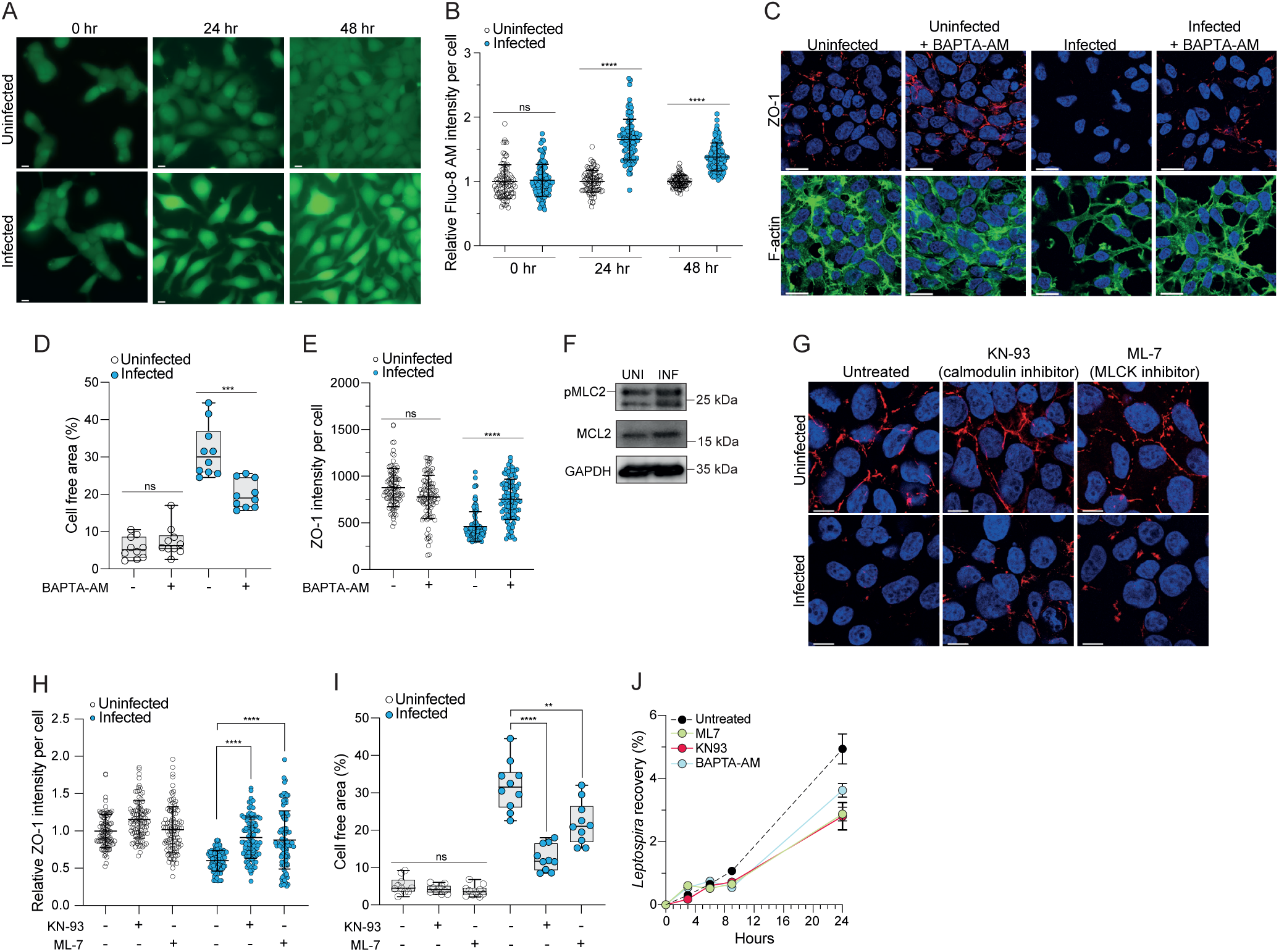
Disruption of cell-cell junctions is partially induced by the calcium signaling pathway through calmodulin and MLCK activities during *L. interrogans* infection. (A) Live images of HEK293T cells infected (lower panels) or not (upper panels) with *L. interrogans* for 0, 24 or 48 hr and stained with Fluo-8 AM (in green) to visualize free Ca^2+^. Scale bars, 10 µm. The images are representative of 3 biological replicates. (B) Quantification of the Fluo-8 AM intensity per cell at 0, 24 and 48 hr pi (n = 100 cells per condition). Values obtained for infected cells (blue symbols) were normalized to that of uninfected cells (empty symbols) at 0 hr. ****, *p* < 0.0001 (unpaired, two-tailed *t* test). (C) Confocal images of human epithelial cells infected or not with *L. interrogans*, treated with the intracellular calcium chelator BAPTA-AM (10 µM) during 24 hr and stained for F-actin (in green, lower panels), ZO-1 (in red, upper panels) and DNA (in blue). Scale bars, 10 µm. The images are representative of 3 biological replicates. (D) Quantification of the percentage of free cell area for uninfected (empty symbols) and infected cells (blue symbols) treated or not with BAPTA-AM as performed in Fig. 5C (n=10 fields of view). Box and Whisker plots represent the median and the interquartile range ± min and max values. Data are representative of 3 biological replicates. ***, p < 0.001 (unpaired, two-tailed *t* test). (E) Quantification of ZO-1 intensity per cell of uninfected (empty symbols) and *L. interrogans*-infected (blue symbols) cells, treated or not with BAPTA-AM as performed in Fig. 5C (n = 100 cells per condition). ****, *p* < 0.0001 (unpaired, two-tailed *t* test). (F) Immunoblot detection of pMyosin (pMLC2), Myosin (MLC2) and GAPDH in total lysates of infected cells (INF) or not (UNI) with *L. interrogans* for 24 hr. GAPDH cellular content was used as a control of equal loading. The molecular mass markers (in kilodaltons) are indicated on the right. The data are representative of 2 biological replicates. (G) Confocal images of uninfected (upper panels) or *L. interrogans*-infected (lower panels) cells treated or not with a calmodulin inhibitor (KN-93, 2 µM) or with a MLCK inhibitor (ML-7, 10 µM) during 24 hr. Cells were stained for ZO-1 (in red) and DNA (in blue). Scale bars, 20 µm. The images are representative of 3 biological replicates. (H) Quantification of ZO-1 intensity per cell of uninfected (empty symbols) or *L. interrogans*-infected (blue symbols) cells treated or not with KN-93 or ML-7 as performed in Fig. 5G. Values were normalized to the uninfected untreated condition (n = 100 cells per condition). The data are representative of 3 biological replicates. ****, *p* < 0.0001 (unpaired, two-tailed *t* test). (I) Quantification of the percent cell-free area for uninfected (empty symbols) or infected (blue symbols) cells treated with KN-93 or ML-7 as performed in Fig. 5G (n=10 fields of view). Box and Whisker plots represent the median and the interquartile range ± min and max values. The data are representative of 3 biological replicates. **, *p* < 0.002 ****; *p* < 0.0001 (unpaired, two-tailed *t* test). (J) Ability of *L. interrogans* to translocate through human epithelial HEK293T cells treated or not (black circles) with BAPTA-AM (blue circles), KN-93 (red circles) or ML-7 (green circles) inhibitors. Bacteria were inoculated in the upper chamber of a cell culture insert containing human epithelial cell monolayer. The percentage of *Leptospira* recovery was determined as described in Fig 4L. Data are mean ± SD from 3 independent experiments.

TJ integrity depends on the interaction between the TJ protein complexes and the perijunctional actomyosin ring^13^, a process controlled by calmodulin-dependent myosin light chain kinase (MLCK) activity. MLCK activation leads to myosin phosphorylation, resulting in increased TJ permeability in epithelial cells^14,15^. *L. interrogans*-infected epithelial cells exhibited elevated level of phosphorylated myosin compared to uninfected controls (**Fig. 5F**). Inhibition of calmodulin or MLCK using KN-93 or ML-7 inhibitors, respectively, partially restored ZO-1 localization at cell-cell junctions and reduced cell dispersion in infected cells (**Fig. 5G-I)**. Notably, this inhibition also impaired *L. interrogans* transmigration across the epithelial monolayer (**Fig. 5J**). These findings indicate that *L. interrogans* exploits host Ca^2+^ signaling to activate the calmodulin-MLCK pathway, leading to junctional disassembly and enhanced paracellular transmigration.

### Two *Leptospira* Virulence-Modifying (VM) proteins are required for virulence and disruption of cell-cell junction through calcium signaling pathway

The most highly upregulated leptospiral genes *in vivo* encode two Virulence-Modifying (VM) proteins (*LIMLP_11655*, Log2FC=7.9; *LIMLP_11660*, Log2FC=8.9; **Source Data Fig.3**). These VM proteins belong to the DUF1561 family, whose function remains unknown. They possess toxin-like features, including a ricin B-like (or Cards B-like) domain potentially mediating interaction with host receptors, and a C-terminal domain of unknown function^16^ (**Extended data Fig. 6A)**. In *L. interrogans*, *LIMLP_11655* and *LIMLP_11660* genes are adjacent (**Extended data Fig. 6B).**

Orthologs of VMs are present in other pathogenic bacteria such as *Bartonella*, *Helicobacter* and *Campylobacter* (**Extended data Fig. 6C-E**). *L. interrogans* harbors a total of thirteen VM proteins, all exclusive to pathogenic *Leptospira* species^8^. Comparative sequence identity analyses indicated that LIMLP_11655 and LIMLP_11660 exhibit greater sequence similarity to each other than to other VM proteins (**Extended data Fig. 6F)**. Notably, LIMLP_11655 and LIMLP_11660 were the only two VM genes significantly upregulated *in vivo* (**Extended data Fig. 6F**).

We had already demonstrated that silencing *LIMLP_11655* (hereafter referred to as *dcas9-LIMLP_11655* mutant) led to an attenuated *L. interrogans* virulence in hamsters^8^. Likewise, *LIMLP_11660* inactivation led to complete loss of virulence, which was restored upon complementation (**Fig. 6A**). Furthermore, when both VM genes were inactivated or silenced, (hereafter referred to as *LIMLP_11660+dcas9-LIMLP_11655 mutant*), *L. interrogans* virulence was lost. Importantly, all these strains exhibited growth rates comparable to that of the WT when cultivated in EMJH medium (**Extended Data Fig. 7**), ensuring that the observed decreased virulence could not be attributed to growth defects. Additionally, all three mutants (*LIMLP_11660*, *dcas9-LIMLP_11655* and *LIMLP_11660+dcas9-LIMLP_11655*) exhibited a reduced ability to transmigrate across epithelial cells at 3 hr pi (**Fig. 6B**).

**Figure 6.**
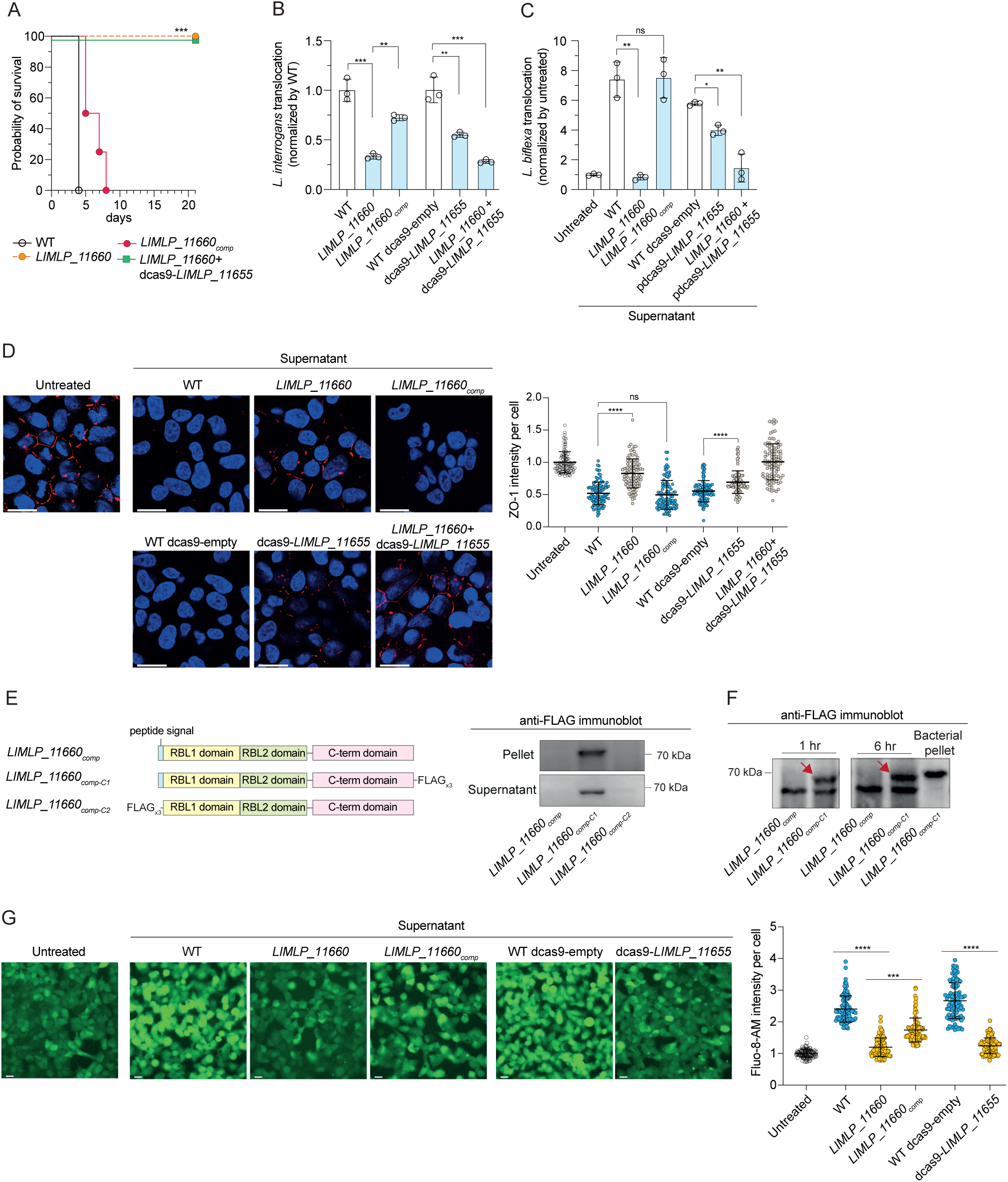
Disruption of cell-cell junctions is mediated by the secretion and internalization of leptospiral virulence-modifying proteins via calcium homeostasis regulation. (B) Survival of hamsters infected with *L. interrogans* WT (empty circles), *LIMLP_11660* mutant (*LIMLP_*11660, orange circles), complemented *LIMLP_11660* mutant (*LIMLP_11660*comp, red circles) or *LIMLP_11660+dcas9-LIMLP_11655* mutant strains (green squares). Groups of 4 animals were infected intraperitoneally with 10^8^ leptospires as described in the Methods section. The WT and *LIMLP_11660* mutant strains contain the empty pMaORI vector. Statistical significance in comparison with the WT strain was determined by a Log rank Mantel Cox test (***, *p* <0.001). (C) *L. interrogans* strains (as designated in Fig. 6A) were added into the upper chamber of a cell culture insert containing a monolayer of human epithelial cell and recovery of *Leptospira* strains in the lower chamber was determined after 3 hr as described for Fig.4L. Values were normalized by those with the WT strain. The dcas9-LIMLP_11665 is a LIMLP_11665 knock down strain and the WT dcas9-empty strain contains the pMaORI vector with the dcas9 sequence used here as a control for dcas9-LIMLP_11665. Data are mean ± SD from 3 independent experiments. **, *p* < 0.002; ***, *p* < 0.0002 (unpaired, two-tailed *t* test). (D) A monolayer of epithelial cells in the upper chamber of a cell culture insert were first incubated with the supernatants from *L. interrogans* cultures (as designated in Figs. 6A and 6B) for 24 hr. *L. biflexa* was then added into the upper chamber. Recovery of *L. biflexa* in the lower chamber was determined after 3 hr by qPCR analysis of the *flab4* gene. Values were normalized by those obtained without the preincubation with *L. interrogans* supernatant cultures (untreated). *, *p* < 0.02; **, *p* < 0.002 (unpaired, two-tailed *t* test). (E) Confocal images (left panel) of human epithelial cells incubated with the supernatant from *L. interrogans* cultures (strains as described for Figs 6A and 6B) for 24 hr and stained for ZO-1 (in red) and DNA (in blue). Scale bars, 20 µm. The quantification of ZO-1 intensity per cell (n=100 cells) is presented in the panel on the right with data normalized by those from uninfected condition. The data are representative of 3 biological replicates. ****, *p* < 0.0001 (unpaired, two-tailed *t* test). (F) LIMLP_11660 is secreted by *L. interrogans*. Expression constructs used to complement the *LIMLP_11660* mutant are represented on the right with *LIMLP_11660comp-C1* corresponding to a C-terminal 3xFLAG tagged LIMLP_11660, and *LIMLP_11660comp-C2* corresponding to a N-terminal 3xFLAG tagged LIMLP_11660 that does not contain any peptide signal. RBL refers to the ricin-B like domain. The presence of FLAG-tagged LIMLP_11660 (expressed *in trans* in the *L. interrogans LIMLP_11660* mutant) in the pellet or supernatant has been evaluated by an anti-FLAG immunoblot (in the right panel). The molecular mass markers (in kilodaltons) are indicated on the right. The data are representative of 3 biological replicates. (G) LIMLP_11660 is internalized inside host cells. Immunoblot detection of FLAG-tagged LIMLP_11660 in the total lysates of host cells incubated with the supernatants of *L. interrogans* cultures (as described above) for 1-and 6 hr. The immunoblot detection of LIMLP_11660comp-C1 in the *L. interrogans* pellet is used here as a positive control. The red arrows indicate the band corresponding to the FLAG-tagged LIMLP_11660. Note that the lower band corresponds to a non-specific recognition. The molecular mass markers (in kilodaltons) are indicated on the left. The data are representative of 3 biological replicates. (H) Live images of epithelial cells (left panel) incubated with the supernatants from *L. interrogans* cultures (strains as described above) for 24 hr and stained with Fluo-8 AM (in green) to visualize free Ca^2+^. Scale bars, 10 µm. The quantification of the Fluo-8 AM intensity per cell (n=100 cells per condition) is presented in the panel on the right with values obtained for infected cells (blue and yellow symbols) normalized to that of uninfected cells (empty symbols). The data are representative of 3 biological replicates. ***, *p* < 0.0002; ****, *p* < 0.0001 (unpaired, two-tailed *t* test).

To investigate whether LIMLP_11660 and LIMLP_11655 contribute to epithelial cell barrier disruption as secreted factors, we tested whether supernatants from WT and mutant strains (*LIMLP_11660*, dcas9-*LIMLP_11655*, and *LIMLP_11660*+dcas9-*LIMLP_11655*) could promote the transmigration of saprophytic *L. biflexa*, normally unable of crossing the epithelial barrier (as seen in **Fig. 4L**). Incubation of epithelial cells with *L. interrogans* WT supernatant enhanced *L. biflexa* translocation, indicating that *L. interrogans* secretes factors capable of disrupting epithelial junctions (**Fig. 6C**). Conversely, supernatants from VM mutants (*LIMLP_11660*, *dcas9-LIMLP_11655* and *LIMLP_11660+dcas9-LIMLP_11655*) exhibited a lower ability to promote *L. biflexa* transmigration and to reduce ZO-1 levels at epithelial junctions (**Fig. 6C-D**). Complementation of *LIMLP_11660* mutant restored both the ability to promote *L. biflexa* transmigration as well as reduced ZO-1 amounts at the cell-cell junction to a comparable manner as the WT (**Fig. 6C-D**).

To confirm VM secretion, we complemented the *LIMLP_11660* mutant with a Flag-tagged allele (*LIMLP_11660comp-C1*) and we successfully detected the Flag-tagged LIMLP_11660 protein in the *L. interrogans* supernatant (**Fig. 6E**). Furthermore, when epithelial cells were exposed to the supernatant from *LIMLP_11660comp-C1* strain for 1 and 6 hr, the Flag-tagged protein was detected in a total lysate of the epithelial cells, and this correlated with a reduced ZO-1 amount at the cell-cell junction comparable to that observed upon infection with the WT strain (**Fig. 6F, Extended Data Fig. 8**). These findings demonstrate that LIMLP_11660-encoded VM is both secreted by *L. interrogans* and internalized by host cells. These results suggest that these two VM proteins exert their role in *Leptospira* pathogenicity by being secreted, binding to the host cell plasma membrane and internalization, disrupting TJ, promoting *Leptospira* transmigrating through epithelial cells layers.

We next examined whether LIMLP_11660 and LIMLP_11655 promoted TJ disruption by altering Ca^2+^-calmodulin-MLCK signaling. Infection with *LIMLP_11660* or *dcas9-LIMLP_11655* mutants significantly impaired the rise of intracellular Ca^2+^ concentration by 4 and 8.5 times, respectively, compared to WT infections (**Fig. 6G)**, and *LIMLP_11660* complementation restored the increase in intracellular Ca^2+^ concentration observed upon infection with the WT strain. This defect correlated with reduced phosphorylation of myosin, a mediator of TJ integrity (**Extended Data Fig. 9)**. Collectively, these findings demonstrate that LIMLP_11660 and LIMLP_11655 act upstream of the Ca^2+^-calmodulin-MLCK pathway, facilitating TJ disruption through calcium signaling activation.

## DISCUSSION

Dual RNA-seq has emerged as a powerful tool to unravel the complex interactions between a host and its pathogen(s), enabling comprehensive understanding of essential biological processes in both organisms during infection. However, its *in vivo* application presents significant technical challenges due to the low bacterial/host RNA ratio, particularly with strict extracellular pathogens where specific selection of infected cells cannot be used. Through advances in high-throughput sequencing technologies, we overcame these obstacles. A deep sequencing approach allowed obtaining robust dual RNA-seq datasets from infected hamsters. This strategy enhanced the coverage of *L. interrogans* transcripts without introducing external factors that might interfere with the host-pathogen dynamics. This refined approach led to valuable insights into the interaction between a host and an extracellular pathogen, providing a framework for studying the molecular mechanisms that drive acute bacterial infection *in vivo*.

The present study demonstrates that, despite *Leptospira* colonizing organs within the first hour of infection^17^, a low transcriptional host response is observed during the first 24h of infection. This lack of early response is particularly striking when compared to other bacterial infections, where early transcriptional responses encompasses about 1,971 DEGs in average across diverse pathogens, regardless of their extracellular or intracellular nature^18–23^ (**Supplementary Table 1**). The low transcriptional host response at the early stage of infection observed here is consistent with the stealth strategy used by pathogenic *Leptospira* to evade host recognition, a key process that favors bacterial dissemination. Indeed, unlike most pathogens, which promptly elicit a strong inflammatory response, *Leptospira* can evade complement-mediated killing^24^, recognition by Toll-like receptor^25^, and internalization by macrophages^8,26^. However, this initial immune evasion is eventually counteracted as the infection progresses. At 3 dpi, the host transcriptomic profile demonstrated a significant upregulation of inflammatory mediators, notably cytokine-cytokine receptor interactions and NF-κB signaling pathways. Among these, genes such as *ccl3*, *cxcl10*, *ccl2*, *tnf*, and *il1β* were highly induced, aligning with previous reports that have implicated these cytokines as potential biomarkers for severe leptospirosis in patients^27–29^. Importantly, our analysis also revealed the upregulation of chemokines and cytokines not previously associated with *L. interrogans* infection, including GDG15 (Growth differentiation factor 15) and CCL4 (also called MIP-1β), which are known to be induced during systemic bacterial infections and sepsis^30,31^. Identifying such biomarkers not only enhances our understanding of the molecular mechanisms of bacterial infection but may also improve diagnosis, which remains challenging, and patient outcomes of this treatable infection.

The liver and the kidney exhibited distinct transcriptomic responses to *L. interrogans* infection, highlighting the importance of organ-specific immune and metabolic adaptations during bacterial colonization. The liver showed a strong upregulation of inflammatory pathways, calcium signaling and cell junction-related genes, probably due to a higher bacterial load (**Fig. 1B**) and extensive vascularization. If so, organ perfusion prior to RNA isolation would allow eliminating such bias. Nevertheless, the attenuated response of kidney observed here could reflect the limited immune cell infiltration of renal proximal tubules, a primary site of *Leptospira* persistence. This strategy would be similar to pathogens, such as *Mycobacterium tuberculosis*^32^ or *Brucella*^33^, that exploit immune-privileged niches to establish long-term colonization.

Breaking the epithelial barrier is a critical step in bacterial pathogenesis but the underlying host-pathogen interactions remain poorly understood, as studies often focus on bacterial factors or the downstream consequences of barrier disruption rather than the molecular mechanisms driving these processes^34,35^. Here we found that several genes related to actin and myosin cytoskeleton, tight and adherens junctions and cell adhesion were upregulated upon infection by *Leptospira*, which is consistent with cell-cell disruption favoring dissemination by a paracellular route^5,36^. Such gene regulations were not observed upon infection by *Salmonella enterica* and *Listeria monocytogenes*, although these pathogens use the paracellular route^19,37^. However, the absence of regulation in cell-cell junction pathway in these bacteria is likely due to their ability to exploit additional mechanisms for host invasion and dissemination. This probably reflects the importance of paracellular route in leptospiral infection.

We demonstrate here that, upon infection by *Leptospira*, modulation of expression of genes related to the transmigration pathway correlated with alterations in intracellular calcium concentration and activation of the MLCK-calmodulin pathway, two processes participating in disruption of cellular junctions. These findings align with a recent study suggesting that the ROCK protein plays a role in E-cadherin disruption in endothelial cells, where ROCK, like MLCK, could induce myosin phosphorylation^36,38^.

Importantly, we could identify in the present study two proteins from the VM family, LIMLP_11660 and LIMLP_11655, as key effectors promoting bacterial dissemination through activation of the Ca^2+^-MLCK-Myosin pathway-induced cell-cell junction disruption. VM proteins exhibit structural features characteristic of AB toxin, a diverse class of bacterial virulence factors. The VMs family comprises 13 members in *Leptospira*, but their specific roles in pathogenicity remain unknown. It was previously showed that all VM genes were upregulated in the blood and liver of infected hamster at 4 dpi, but different expression levels were observed in the kidneys^40^. The two VMs studied here were the most highly expressed of the VMs in the condition used here. This suggests not only a temporal and tissue-specific expression pattern but also distinct function in infection. Even though some VMs may exhibit redundant activity, such as the two VMs identified here, others very likely contribute to different aspects of *Leptospira* pathogenesis, such as immune evasion, adhesion to host tissues, modulation of host signaling pathways or/and persistence in specific organs. Further functional characterization is required to elucidate the precise contributions of each VM in *Leptospira* virulence and dissemination.

Unlike most pathogenic diderm bacteria which encode type III and IV secretion systems to directly translocate effector proteins into the cytoplasm or plasma membrane of host cells^41^, *Leptospira* encode only type I and II secretion systems. These systems are responsible for exporting proteins into the extracellular environment, suggesting a distinct strategy for host-pathogen interactions. Interestingly, leptospiral type I and II secretion systems were upregulated upon infection (as demonstrated in this study), suggesting a general role during infection and perhaps particularly in the secretion of virulence factors such as VMs.

By linking host calcium signaling and leptospiral VM-induced TJ disruption, we provide here a novel mechanism for bacterial pathogenesis. We propose that, once within the host, *Leptospira* induce the expression and secretion of these two VMs. These proteins interact with epithelial cells likely by binding of their Ricin B domain to lectin receptors. The VMs would be then internalized inside host cells, and they would modulate calcium homeostasis, either directly or indirectly. This would result in the activation of the calmodulin-MLCK signaling pathway, ultimately leading to TJs disassembly (**Supplementary Fig. 2**). The loss of barrier integrity facilitates the translocation of *Leptospira* through epithelial and endothelial layers, promoting systemic dissemination to target organs. Interestingly, VM proteins are not exclusive to pathogenic *Leptospira* but are also found in other pathogenic bacteria, including *H. pylori*, *C. jejuni* and *Bartonella*, which are known to target host cell-cell junctions for bacterial dissemination. VMs have never been studied nor characterized in these bacteria but they may function similarly to those in *L. interrogans*, by disrupting epithelial barriers through a host calcium homeostasis-controlled mechanism. In fact, this novel mechanism of cell-cell junction disruption may not be exclusive to *Leptospira* and could also be exploited by other pathogens, whether of bacterial, fungal or viral origin, to facilitate their dissemination across host barriers.

## METHODS

### Bacterial strains and culture conditions

Strains and plasmids used in this study are listed in **Supplementary Tables 2 and 3**. WT *L. interrogans* serovar Manilae strain L495 and mutants were cultivated aerobically in Ellighausen-McCullough-Johnson-Harris liquid medium (EMJH) at 30°C with shaking at 100 rpm. ρχ1 and β2163 *E. coli* strains were cultivated at 37°C in Luria-Bertani medium with shaking at 37°C in the presence of 0.3 mM thymidine or diaminopimelic acid (Sigma-Aldrich), respectively. Spectinomycin was added to the media at 50 µg/ml when needed.

### *In vivo* animal infection

Four-week-old male Golden Syrian hamsters (RjHan:AURA, Janvier Labs) (n=4) were infected via intraperitoneal injection with 10^8^ *L. interrogans* WT and mutant strains, with the number of bacteria enumerated using a Petroff-Hausser counting chamber. The animals were monitored daily and euthanized by carbon dioxide inhalation when they reached the predefined endpoint criteria (sign of distress). To assess leptospiral load, samples of kidney and liver tissue were harvested, and DNA was extracted using the Tissue DNA Purification kit (Maxwell, Promega). The bacterial burden and host DNA concentration were determined by quantitative polymerase chain reaction (qPCR) using the Sso Fast EvaGreen Supermix assay (Bio-Rad) with primers targeting the *flaB2* (LIMLP_09410) and *gapdh* genes, respectively. The leptospiral load was expressed as genomic equivalents (GEq) per microgram of host DNA.

### RNA extraction, library preparation and sequencing

Liver and kidneys tissues of hamsters (infected or not by *L. interrogans* as described above) were resuspended in the QIAzol lysis reagent (Qiagen) with 5 mm stainless-steel beads (Qiagen) and homogenized with the Qiagen TissueLyser (two cycles of one-minute at 25 Hz, with a one-minute break between cycles in ice). Total RNAs were extracted from homogenized samples using the RNeasy Mini Kit (Qiagen), with a DNAse digestion (Qiagen) incorporated in the protocol. RNAs were also isolated from exponentially growing *L. interrogans* cultivated in EMJH medium. Bacterial pellets were resuspended in the QIAzol lysis reagent and total RNAs were extracted using a similar procedure (RNeasy Mini Kit with a DNAse treatment). The quality of all RNA samples was evaluated using an Agilent 2100 bioanalyzer (Agilent Technologies) to ascertain a RNA integrity number (RIN) higher than 8.5.

Sequencing Libraries were constructed using an Illumina Stranded Total RNA Prep Ligation using the RibZero Plus kit (Illumina, USA) with custom primer design on rRNA following the supplier’s recommendations. RNA sequencing was performed with the Illumina NextSeq 2000 and NovaSeq X+ platform for a target of 10M reads/sample for bacteria, 30M reads/sample for hamster and 400M reads/sample for deep sequencing (performed on RNA samples isolated from 3 hamsters infected by *L. interrogans*). The RNA-seq analysis was performed with Sequana 0.15.3^42^ using the RNA-seq pipeline 0.16 (https://github.com/sequana/sequana_rnaseq) built on top of Snakemake 7.32.4^43^. Briefly, reads were trimmed from adapters using Fastp 0.22.0^44^ then mapped to the *L. interrogans* serovar Manilae genome (NZ_CP011931.1) and Golden Syrian hamsters genome (KB708127.1) using Bowtie2^45^ and STAR^46^. FeatureCounts 2.0.1^47^ was used to produce the count matrix, assigning reads to features using the corresponding annotation from Ensembl with strand-specificity information. Quality control statistics were summarized using MultiQC 1.11^48^. Statistical analysis on the count matrix was performed to identify differentially regulated genes. Clustering of transcriptomic profiles were assessed using a Principal Component Analysis (PCA). Differential expression testing was conducted using DESeq2 library 1.34.0^49^ scripts indicating the significance (Benjamini-Hochberg adjusted p-values, false discovery rate FDR < 0.05) and the effect size (fold-change) for each comparison.

Gene ontology (GO) enrichment analyses were performed using the Cytoscape app ClueGO (version 2.5.3)^50^, with the following parameters: only pathways with pV ≤0.01, Minimum GO level = 3, Maximum GO level = 8, Min GO family >1, minimum number of genes associated to GO term = 5, and minimum percentage of genes associated to GO term = 8. Enrichment p-values were calculated using a hypergeometric test (p-value<0.05, Bonferroni corrected).

### Gene expression by RT-qPCR

Reverse transcription of mRNA to cDNA was performed using iScript cDNA Synthesis kit (Bio-Rad), followed by cDNA amplification using the SsoFast EvaGreen Supermix (Bio-Rad). All primers used in this study are listed in **Supplementary Table 4**. Reactions were performed using the CFX96 real-time PCR detection system (Bio-Rad). The relative gene expression levels were assessed according to the 2^−ΔΔCt^ method using *flab2* or *gapdh* as reference gene for *L. interrogans* or human cells, respectively.

### Cell culture and infection

HEK-293T cells were cultured in Minimum Essential Medium Eagle (MEME) (Sigma) supplemented with 10% heat-inactivated fetal bovine serum (Sigma) and 2 mM L-glutamine (Gibco). *Leptospira* strains were diluted in cell culture media prior the infection, at a multiplicity of infection of 100:1. All chemical compounds used on epithelial cells were added 1 hr before the experiment and maintained throughout the duration specified in the figure legend.

### Indirect immunofluorescence and live-cell imaging of epithelial cells

Epithelial cells were cultured on glass coverslips (SPL) coated with L-lysine (Sigma) at a concentration of 0.01% in water for a period of 40 min at 37°C. The coverslips were rinsed twice with PBS to remove excess of L-lysine, and the cells were seeded. At the indicated times, the cells were fixed with 4% paraformaldehyde for 15 min at room temperature (RT) and subsequently incubated for 10 min in 0.5% saponin (Sigma) in PBS and for 1 hr in 1% BSA (Sigma) and 0.075% saponin in PBS. The cells were incubated overnight at 4°C with the anti-ZO1 (13663; Cell Signaling), anti-ZO2 (2847; Cell Signaling), anti-F-actin (PHDH1; Tebu), anti-JAM1 (82196; Cell Signaling) or anti-connexin-43 (3512; Cell Signaling) antibodies at 1:100. Cells were washed and incubated for 1 hr with Alexa Fluor 555 or 488 antibody (ThermoFisher) at 1:500. The nuclei were then stained with DAPI (1 µg/mL; ThermoFisher) for 10 min and mounted on a glass side using Fluoromount mounting medium (ThermoFisher). Fluorescence was analyzed using a Leica TCS SP8 Confocal System, and the quantification was performed using Icy software.

For live-cell imaging, cells were seeded on an µ-Slide 8 Well (Ibidi). To assess ER activity, cells were labelled with ER-Tracker Red (ThermoFisher) at a concentration of 1 µM for 30 min at 37°C, followed by two washes with FluoroBrite DMEM medium (ThermoFisher) supplemented with 10% heat-inactivated fetal bovine serum (Sigma) and 2 mM L-glutamine (Gibco). For the measurement of intracellular calcium levels, cells were incubated with Fluo-8 AM (Abcam) at a concentration of 4 µM for 1 hr, after which they were washed twice with complete FluoroBrite DMEM medium prior to the analysis time. Brightfield and fluorescence microscopy were conducted using an inverted widefield microscope equipped with an LED colibris and a mercury lamp. Images were acquired using a dual camera sCMOS Hamamatsu ORCA FLASH. Cell dispersion and fluorescence quantification were performed using Icy software.

### Cell viability assay

Cell viability was determined using the MTT assay kit (Abcam), according to manufacturer’s instructions.

### Western Blot assay

Total extracts of epithelial cells or *Leptospira* strains were obtained by sonication in a lysis buffer containing 25 mM Tris pH 7.5, 100 mM KCl, 2 mM EDTA, 5 mM DTT and a protease inhibitor cocktails (cOmplete Mini EDTA-free, Roche). 10 µg of total proteins were loaded on a 12% SDS-PAGE and transferred onto nitrocellulose membrane. The membranes were blocked with PBS Tween20 0.1% containing 5% BSA for 1 hr at RT and incubated overnight at 4°C with the antibodies anti-ZO1 (13663; Cell Signaling), anti-ZO2 (2847; Cell Signaling), anti-Profilin-1 (3246; Cell Signaling), anti-WAVE-2 (3659; Cell Signaling), anti-pRac1 (2461; Cell Signaling), anti-pCofilin (3313; Cell Signaling), anti-Cofilin (5175; Cell Signaling), anti-GAPDH (5174; Cell Signaling), anti-MLC2 (8505; Cell Signaling), anti-pMLC2 (3671; Cell Signaling), anti-PERK (3192; Cell Signaling), anti-pPERK (PA5-40294; ThermoFisher), anti-IRE1 (3294; Cell Signaling), anti-pIRE1 (PA1-16927; ThermoFisher) and anti-3xFLAG (87537; Cell Signaling) at a 1:1000 dilution. Membranes were washed in PBS-0.1% Tween20 and incubated at RT with secondary HRP-conjugated antibody (7074S; Cell Signaling) at a 1:1000 dilution for 1 hr. Detection was performed with SuperSignal West Femto Maximum Sensitivity Substrate (Thermo Fisher). Quantification of band intensities was performed using ImageJ software.

### Translocation through epithelial cells

5.10^4^ HEK293T cells in 300 µL of complete MEME were introduced into 12-mm-diameter Transwell filter units with 3 µm pores (COSTAR). The monolayers were incubated at 37°C in 5% CO2 for a period of three days until the transepithelial resistance reached a range of 200 and 300 Ω/cm2 using an epithelial voltammeter. The monolayers were infected with leptospires at a MOI of 100. At the indicated times, the capacity of *Leptospira* strains to translocate across the epithelial cell barrier was evaluated by extracting the DNA from the lower chamber (300 µL) using the Maxwell RSC Cultured Cells DNA Kit and the Maxwell instrument (Promega). The concentration of leptospires was determined by qPCR with the Sso Fast EvaGreen Supermix assay (Bio-Rad) using *flaB2* (LIMLP_09410) for *L. interrogans* and *flaB4* (LEPBIa1589) for *L. biflexa*. The capacity of leptospires to translocate HEKT293T polarized monolayers was quantified by calculating the proportion of leptospires present in the lower chamber relative to the initial inoculum at each time points.

### Complementation of *L. interrogans* mutants

Complementation of the *LIMLP_11660* mutant was performed by expressing LIMLP_11660 using the replicative vector pMaOri^51^ (**Supplementary Table 3)** under the control of its native promoter. For this, LIMLP_11660 ORF and the 300 bp upstream region were amplified from genomic DNA of *L. interrogans* (using CompLIMLP_11660 primer set; **Supplementary Table 4)** and cloned between the SacI and NotI restriction sites in the pMaOri vector. The absence of mutations in the resulting plasmid (pMaOri-*LIMLP_11660*, **Supplementary Table 3**) was confirmed by DNA sequencing (Eurofins). Then, the plasmid pMaOri-*LIMLP_11660* was introduced into *LIMLP_11660* mutant strain by conjugation using the *E. coli* β2163 conjugating strain, as previously described^51^. *Leptospira* conjugants were selected on EMJH agar plates containing 50 µg/mL spectinomycin. WT and *LIMLP_11660* mutant strains containing the empty pMaOri vector were used as controls. Expression of the Flag-tagged LIMLP_11660 variant was performed by introducing a 3xFlag tag in frame at the 3’ of the ORF (*LIMLP_11660comp-C1*) or by replacing the peptide signal (*LIMLP_11660comp-C2*) using restriction-free cloning method in the plasmid pMaOri-*LIMLP_11660* with the primers *LIMLP_11660-C1* and *LIMLP_11660-C2* (**Supplementary Table 4**). Conjugation in *L. interrogans* was performed as described above. Gene silencing of *LIMLP_11655* into the *LIMLP_11660* mutant was performed by introducing the leptospiral replicative vector pMaori.dCas9_sgRNA_*LIMLP_11655* by conjugation as previously described^8^.

### Supernatant isolation of *L. interrogans*

Supernatants were obtained from exponentially growing *L. interrogans* cultivated in Minimum Essential Medium Eagle (MEME, Sigma) supplemented with 10% heat-inactivated fetal bovine serum (Sigma) and 2 mM L-glutamine (Gibco) overnight at 37°C. Bacteria were centrifuged at 5000 g for 15 min and the supernatants were filtered through a 300 kDa membrane (Sartorius).

### *In silico* analysis of LIMLP_11660 and LIMLP_11655

The sequence similarity of the LIMLP_11660 protein from *L. interrogans* serovar Manilae was determined using UniProtKB reference proteomes and Swiss-Prot databases. All significant hits, defined as the best match for each species with an e-value <0.00033 and a protein identity >20%, were retained. Sequence alignment of the proteins and percent identity matrix were performed using Clustal-Omega (1.2.4). Tree inference was achieved with IQ-TREE v2.0.6 under the best-fitted model.

The domain models were created using the LIMLP_11660 or LIMLP_11655 protein sequences via the SWISS-MODEL Workspace server (https://swissmodel.expasy.org/, SMTL version 2023-04-05, PDB release 2023-03-31).

### Ethics Statement

Protocols for animal experiments are conformed to the guidelines of the Animal Care and Use Committees of the Institut Pasteur (Comité d’éthique d’expérimentation animale CETEA # 220016), agreed by the French Ministry of Agriculture. All animal procedures carried out in our study were performed in accordance with the European Union legislation for the protection of animals used for scientific purposes (Directive 2010/63/EU).

### Statistics and reproducibility

Statistical data was conducted using in GraphPad Prism (version 9.5.1). Unless otherwise stated, all experiments were performed in three independent biological replicates. All microscopy images shown are representative of three independent biological replicates. In all graphs, data are presented as mean ± SD. Centre line, median, box limits, the interquartile range ± min and max values were indicated in box and whisker plots. Information about specific statistical tests in each analysis can be found in the figure legends.

## ACKNOWLEDGEMENTS

This work has received financial support by the a PTR (Programmes transversaux de Recherche) grant (PTR2019-310) from Institut Pasteur Paris (to NB), by a Groot-23 from the department of Microbiology of Institut Pasteur Paris (to AGG), and by the National Institutes of Health grant P01 AI 168148 (to MP & NB). We thank the Ultrastructural BioImaging Platform (UTechS UBI, Institut Pasteur Paris) for technical support. We also acknowledge L. Lemée and R. Ouazahrou from Biomics Platform, C2RT, Institut Pasteur, Paris, France, supported by France Génomique (ANR-10-INBS-09) and IBISA for their technical help for RNASeq experiments. We are grateful to N.

Sauvonnet, E. Lemichez, L. Tailleux, and M. Lago for critically reading the manuscript and helpful discussions. The funders had no role in study design, data collection and analysis, decision to publish, or preparation of the manuscript.

