## Supplementary information for "*In Vivo* Dual RNA-Seq uncovers key toxin-like effectors of epithelial barrier disruption and tissue colonization by an extracellular bacterial pathogen"

### EXTENDED DATA

**Extended Data Fig. 1. Proportion of hamster and *L. interrogans* genes detected *in vivo*.**
(**A**) Percentage of reads mapped to the hamster (blue bars) or *L. interrogans* (red bar) genomes for the indicated transcriptome condition. Data are mean and SD from 4 biological replicates per condition. The number of *Leptospira* genes detected are indicated for each condition.
(**B**) Density plot showing the distribution of *L. interrogans* read counts in samples isolated from for *L. interrogans* cultured *in vitro* (red line), in samples from liver of infected hamsters at 3 days post-infection (dpi) sequenced at 40M reads/sample (black line) or at 400M reads/sample (i. e. deep sequencing, yellow line).

**Extended Data Fig. 2. Host transcriptional response to *L. interrogans* infection in liver and kidneys of hamsters.**
**(A)** Venn diagram showing unique and shared DEGs (FDR≤0.05; $|Log_2FC|≥1$) in liver and kidneys at 1- and 3 days post-infection (dpi). Number of up-regulated and down-regulated genes were indicated with a blue and a red arrow, respectively.
**(B)** Gene Ontology (GO) enrichment analysis of upregulated genes (FDR≤0.05) in liver at 3 dpi, using Cytoscape app ClueGO. The biological process and molecular function were used to classify the biological functions of upregulated genes. Each GO term represents a circle with related terms grouped as families represented with a specific color. The lines indicate link between GO terms. Only GO term with Bonferroni adjusted *p*-value<$10^{-4}$ are shown.
**(C)** GO enrichment analysis of downregulated genes (FDR≤0.05) in liver at 3 dpi determined as described in (B).
**(D)** Gene Ontology (GO) enrichment analysis of upregulated genes (FDR≤0.05) in kidneys at 3 dpi, determined as described in (B).

**Extended Data Fig. 3. COG enrichment analysis of *L. interrogans in vivo*.**
Radar graphs representing the clusters of orthologous genes (COGs) analysis of *L. interrogans* DEGs (FDR ≤0.05) at 3 dpi compared to *L. interrogans* cultivated in EMJH medium. COGs categories were divided in three major groups (Cellular processes and signaling, Information storage and processing and Metabolism). Down-regulated ($Log_2FC<0$) and up-regulated ($Log_2FC>0$) ORFs are represented in blue and red, respectively. For each radar axes, the scale corresponding to the number of ORFs in each COG category is indicated.

**Extended Data Fig. 4. *L. biflexa* does not induce actin remodeling or cell-cell junction disruption in human epithelial cells.**

(**A**) Confocal images of human epithelial cells uninfected (upper panel) or infected with *L. biflexa* for 24 hr (lower panel) and stained for F-actin (in green) and DNA (in blue). Scale bars, 10 µm. The images are representative of 3 biological replicates.

(**B**) Quantification of the percentage of cell-free area for uninfected (empty symbols) or infected condition (orange symbols) (n=10 fields of view). Box and whisker plots represent the median and the interquartile range ± min and max values, respectively (unpaired, tailed *t* test).

(**C**) Confocal images of human epithelial cells uninfected (upper panels) or infected with *L. biflexa* for 24 hr (lower panels), stained for ZO-1, ZO-2 or JAM-1 (in red) and DNA (in blue). Scale bars, 20 µm. The images are representative of 3 biological replicates.

(**D**) Fluorescence quantification of ZO-1, ZO-2 and JAM-1 intensity per cell (n=100 cells) in infected (orange symbols) or uninfected (empty symbols) samples. Differences are considered non-significant by the unpaired tailed *t* test.

**Extended Data Fig. 5. Calcium chelation reduces the rise of intracellular calcium in *L. interrogans*-infected cells.**

(**A**) Live images of human epithelial cells untreated (upper panels) or treated with the intracellular calcium chelator BAPTA-AM (10 µM, lower panels), and infected (right panels) or not (left panels) with *L. interrogans* for 24 hr. Cells were stained with Fluo-8 AM (in green) to visualize free $Ca^{2+}$. Scale bars, 20 µm. The images are representative of 3 biological replicates.

(**B**) Quantification of Fluo-8 AM intensity per cell at 24 hr post-infection (n=100 cells per condition) of uninfected (empty symbols) or infected (blue symbols) cells, treated or not with BAPTA-AM as described in (A). ***, $p < 0.0002$ (unpaired, two-tailed *t* test).

**Extended Data Fig. 6. *In silico* characterization of the VM protein family.**

(**A**) Schematic representation of the domain architecture of LIMLP_11660 and LIMLP_11655 (RBL: ricin B-like domain). Domain predictions were obtained using Swiss-model modeling server, with sequence identity percentages shown for each predicted domain.

(**B**) Schematic representation of the genomic localization of LIMLP_11660 and LIMLP_11655 ORFs in the *L. interrogans* UP-MMC-NIID LP strain chromosome (CP011931.1).

(**C**) Phylogenetic tree of LIMLP_11660-related proteins across other bacterial species, constructed using IQ-TREE v2.0.6 under the best-fitted model. Taxonomic groups are shown with different colors. Ortholog sequences for LIMLP_11660 were searched using UniProtKB reference proteomes and Swiss-Prot databases with an e-value <0.00033 and a protein identity > 20%.

(**D**) Matrix displaying the average amino acid identity (AAI) among LIMLP_11660-related proteins with a heat map color from blue to orange indicating low to high % identity.

(**E**) Hosts of bacterial species harboring LIMLP_11660-related proteins.

(**F**) Conservation and *in vivo* expression of all the VM proteins in *L. interrogans*. A phylogenetic tree based on the AAI of the thirteen VM proteins in *L. interrogans* is displayed on the left panel. The matrix displaying the percentage of AAI across VM proteins is shown in the middle panel with a heat map color from blue to orange indicating low to high % identity. The heatmap representation of the *in vivo* expression levels (in $Log_2FC$) of each VM protein is presented in the right panel, with each column corresponding to an independent biological replicate and heat map color from blue to red indicating low to high differential expression. The phylogenetic clustering analysis delineates three distinct VM subgroups.

**Extended Data Fig. 7. Growth of *L. interrogans* strains used in this study**

Growth analysis of *L. interrogans* WT and mutant strains containing different expression constructs. Strains include WT with the empty pMaORI vector (WT, black circles) or an empty pMaORI-dcas9 vector (WT dcas9-empty, blue squares); the *LIMLP_11660* mutant containing the empty pMaORI vector (*LIMLP_11660*, red triangles) or the pMaORI vector expressing *LIMLP_11660* (*LIMLP_11660*$_{comp}$, green inverted triangles); *L. interrogans* WT containing the pMaORI-dcas9 vector expressing the single-guide RNA targeting *LIMLP_11655* (pdcas9-*LIMLP_11655*, yellow diamonds); and the *LIMLP_11660* mutant containing the pMaORI-dcas9 vector expressing the single-guide RNA targeting *LIMLP_11655* (*LIMLP_11660* + pdcas9-*LIMLP_11655*, brown circles). All strains were cultivated in EMJH medium complemented with spectinomycin. Bacterial growth was assessed by measuring of absorbance at 420 nm. Data are the mean ± SD from 3 independent experiments.

**Extended Data Fig. 8. LIMLP_11660 participates in the disruption of cell-cell junction in epithelial cells infected by *L. interrogans*.**

Quantification of ZO-1 intensity per epithelial cell incubated with the supernatant from *L. interrogans* cultures of the WT or the *LIMLP_11660* mutant (expressing different alleles of LIMLP_11660 as described for Fig. 6E) for 24 hr from confocal images (n = 100 cells). Data were normalized with those from uninfected condition. The WT and *LIMLP_11660* mutant strains contain the pMaORI vector. Data are the mean ± SD from triplicates representative of 3 independent biological replicates. **, $p < 0.002$; ***, $p < 0.0002$; ****, $p < 0.0001$ (unpaired, two-tailed *t* test).

**Extended Data Fig. 9. LIMLP_11660 and LIMLP_11655 modulate myosin phosphorylation in human epithelial cells.**

Immunoblot detection (**A**) and quantification (**B**) of phosphorylated (pMLC2) and unphosphorylated myosin (MLC2) proteins in total lysates of cells incubated for 24 hr with the supernatant from *L. interrogans* cultures (as described for Fig. 6). Ratio of phosphorylated over unphosphorylated myosin (pMLC2/MLC2) is shown for each sample. Data are the mean ± SD from 2 biological replicates. *, $p < 0.01$ (unpaired, two-tailed *t* test).

**Extended Data Table 1. Regulated pathways in the liver and kidneys of hamsters infected with *L. interrogans* at 3 days post-infection compared to uninfected controls.**

**Extended Data Table 2. Regulated pathways of *L. interrogans* in the liver of infected hamsters compared to EMJH culture.**

**Extended Data Table 3. Transcriptional regulation of actin cytoskeleton-related genes in the liver and kidneys of *Leptospira*-infected hamsters.**

### SUPPLEMENTARY DATA

**Supplementary Fig. 1. Induction of ER stress response during *L. interrogans* infection is not required for disruption of cell-cell junctions.**

(**A**) Gene Ontology pathways analysis of DEGs related to endoplasmic reticulum activity in the liver of *L. interrogans*-infected hamsters. Only the eight more significative pathways are represented (with a *p*-value corrected with Bonferroni step down < 0.005). The percentage of DEGs associated with each pathway is indicated.

(**B**) Relative expression of *u/sxbp1*, *perk*, *ire1a* and *bax* genes at 24 and 48 hr pi in *Leptospira*-infected epithelial cells compared to uninfected condition. Gene expression levels were determined by RT-qPCR and normalized using *gapdh* gene as reference gene. Data are mean ± SD of 3 biological replicates. *, $p < 0.01$ (unpaired, two-tailed *t* test).

(**C**) Live images of human epithelial cells infected (lower panels) or not (upper panels) with *L. interrogans* for 6, 24 or 48 hr and stained with ER-Tracker (in red) to visualize the ER. Scale bars, 10 µm. The images are representative of 3 biological replicates.

(**D**) Quantification of the ER-Tracker intensity per cell at 6, 24 or 48 hr pi (n=100 cells per condition, in the samples presented in C). ****, $p < 0.0001$ (unpaired, two-tailed *t* test).

(**E**) Immunoblot detection of IRE-1, phosphorylated IRE-1 (pIRE-1), PERK, and phosphorylated (pPERK) in total lysates of HEK293T cells uninfected (UNI) or infected (INF) for 24 and 48 hr with *L. interrogans*. Immunoblots were performed as described in the Methods section. GAPDH cellular content was used as an equal loading control. The molecular mass markers (in kilodaltons) are indicated on the right. Data are representative of 2 independent experiments.

(**F**) Confocal images of human epithelial cells untreated or treated with a PERK inhibitor (GSK2606414, 11 mM) or an IRE-1 inhibitor (STF-083010, 12.5 mM) and infected (lower panels) or not (upper panels) with *L. interrogans* for 24 hr. Cells were stained for ZO-1 (in red) and DNA (in blue). Scale bars, 20 µm. The images are representative of 3 biological replicates.

(**G**) Quantification of ZO-1 intensity per cell (n=100 cells) in the samples presented in (F) and in cells treated with ER stress inhibitors 4-PBA (2mM) or TUDCA (150 µM), infected (blue symbols) or not (empty symbols) with *L. interrogans* for 24 hr. Data were normalized with those from uninfected condition. The data are representative of 3 biological replicates. **** $p < 0.0001$ (unpaired, two-tailed *t* test).

**Supplementary Fig. 2. Proposed model of VM protein-mediated disruption of cell-cell junctions.**
In non-infected cells (on the left), tight junctions (JAM, Occludin and Claudin) stabilize cell-cell junctions and ZO proteins connect tight junctions and actin cytoskeleton. During leptospiral infection (on the right), *L. interrogans* induces the expression and secretion of two virulence-modifying (VM) proteins, LIMLP_11655 and LIMLP_11660. These proteins would interact directly with host epithelial cells, very likely by binding of their Ricin B domain to lectin receptors present at the host cell surface. The VMs are then internalized within the host cell, where they trigger an increase in intracellular calcium levels. This calcium deregulation would activate the calmodulin-MLCK pathway, leading to actin cytoskeleton remodeling and ultimately disassembly of tight junctions. The resulting loss of barrier integrity facilitates *L. interrogans* transmigration across epithelial and endothelial cell layers, promoting systemic dissemination to target organs, such as liver and kidneys.

**Supplementary Table 1. Transcriptional response to infection by different bacterial pathogens.**

**Supplementary Table 2. Bacterial strains used in this study**

**Supplementary Table 3. Plasmids used in this study**

**Supplementary Table 4. Primers used in this study**

### SOURCE DATA

**Source Data Fig. 2.** Differentially expressed genes in the liver and the kidneys of hamsters infected with *L. interrogans* at 1 and 3 day post-infection.
**Source Data Fig. 3.** Differentially expressed genes of *L. interrogans* in the hamster liver at 3 day post-infection compared to *in vitro* condition.

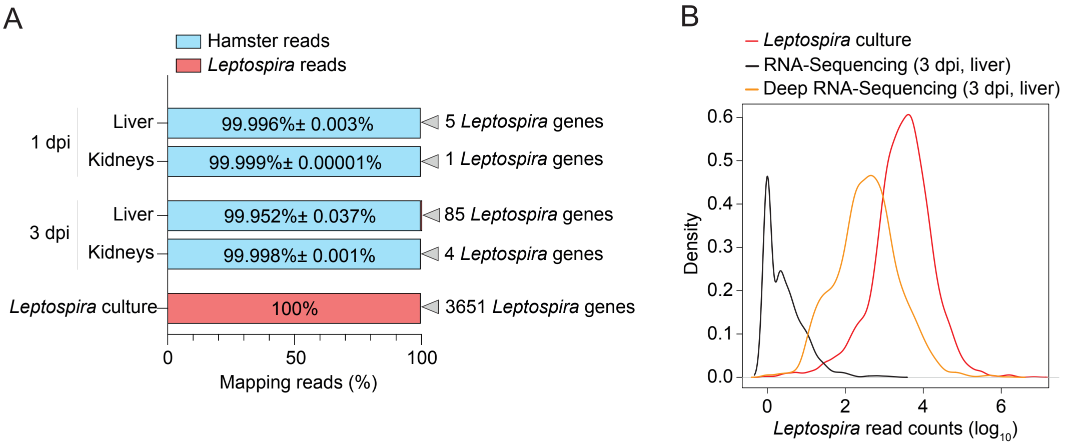


**Extended data Fig. 1**

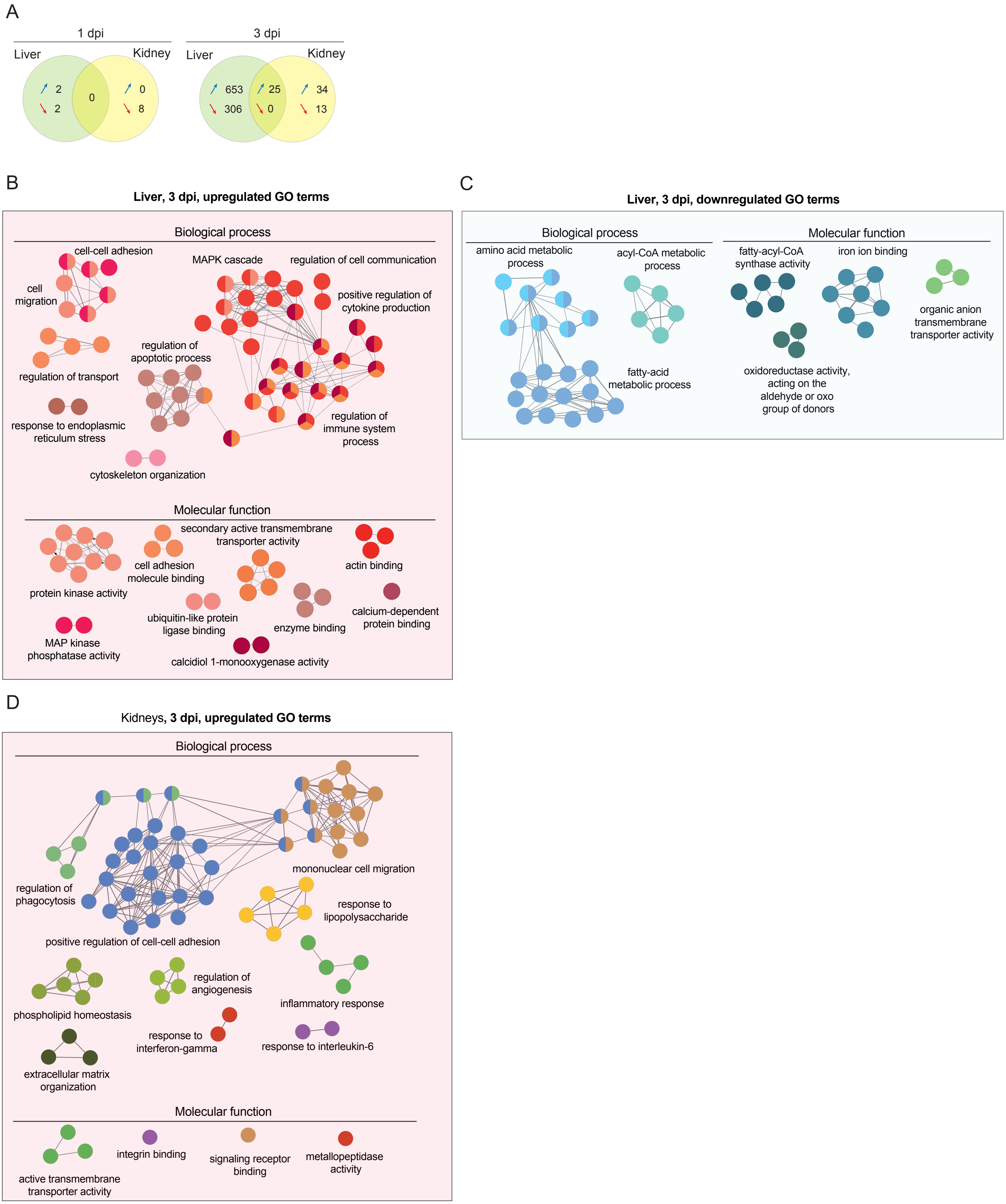


**Extended data Fig. 2**

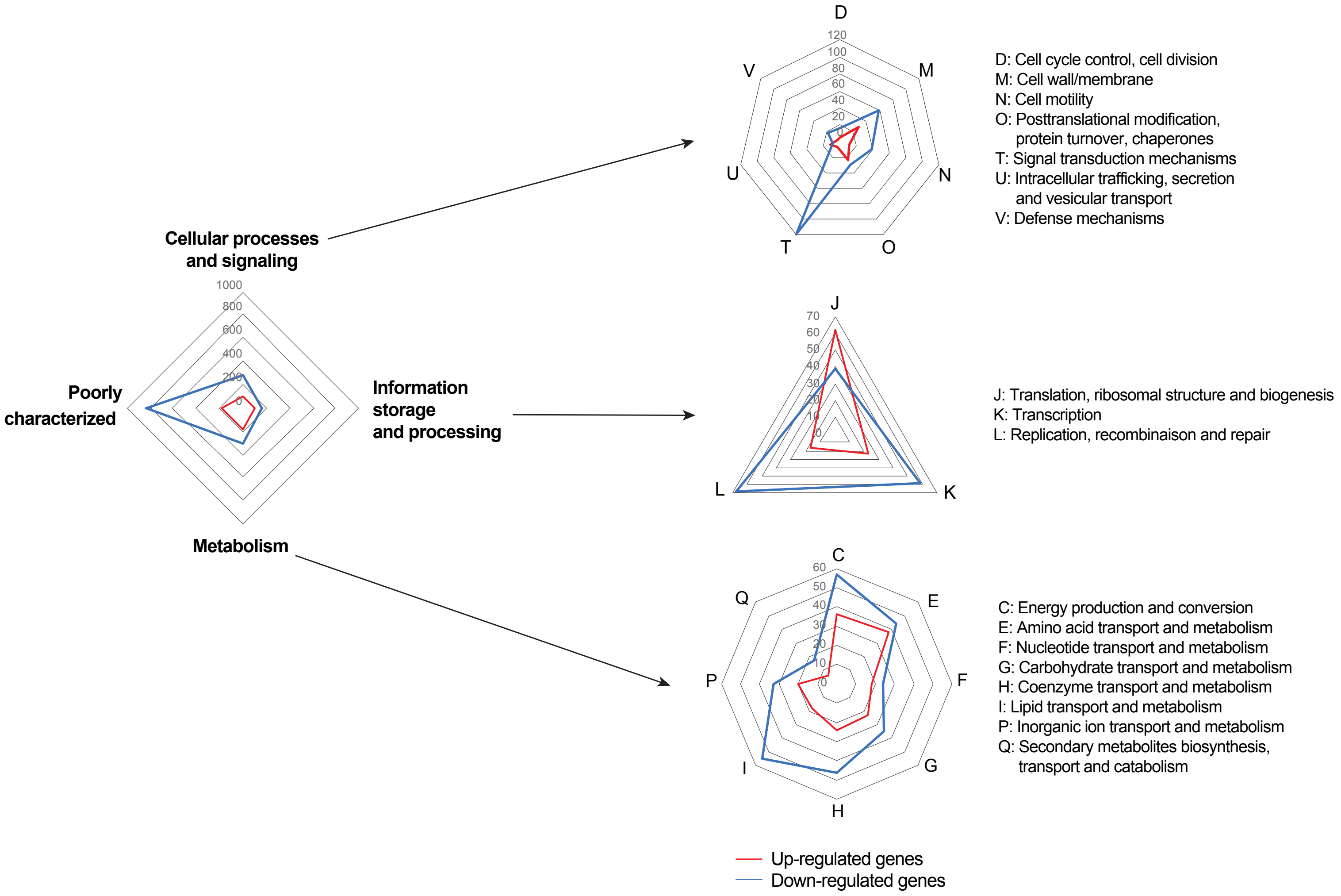


**Extended data Fig. 3**

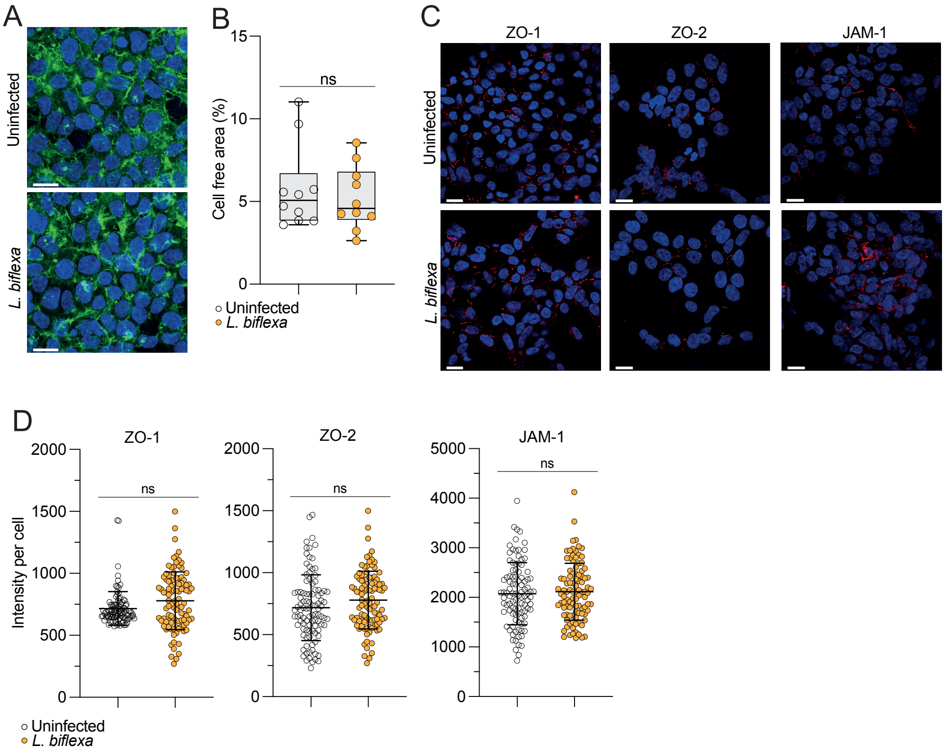


**Extended data Fig. 4**

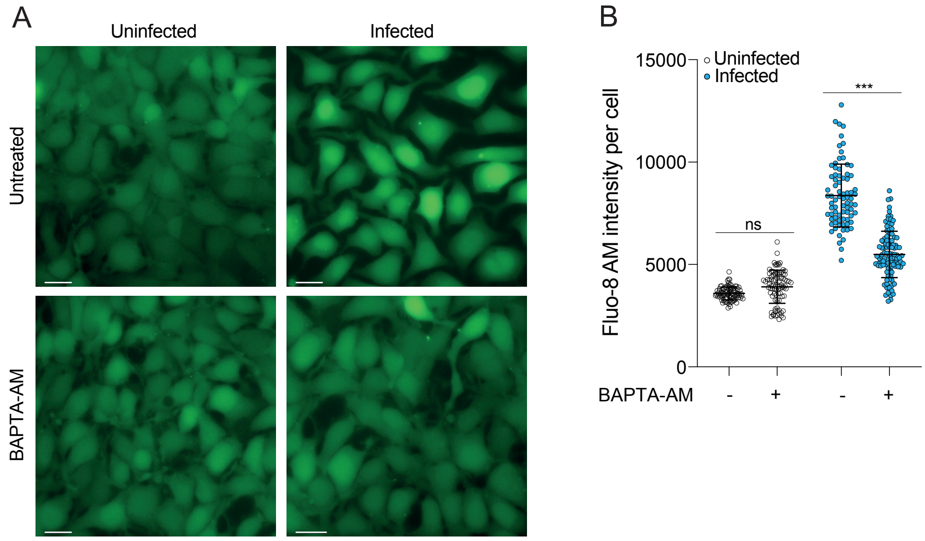


**Extended data Fig. 5**

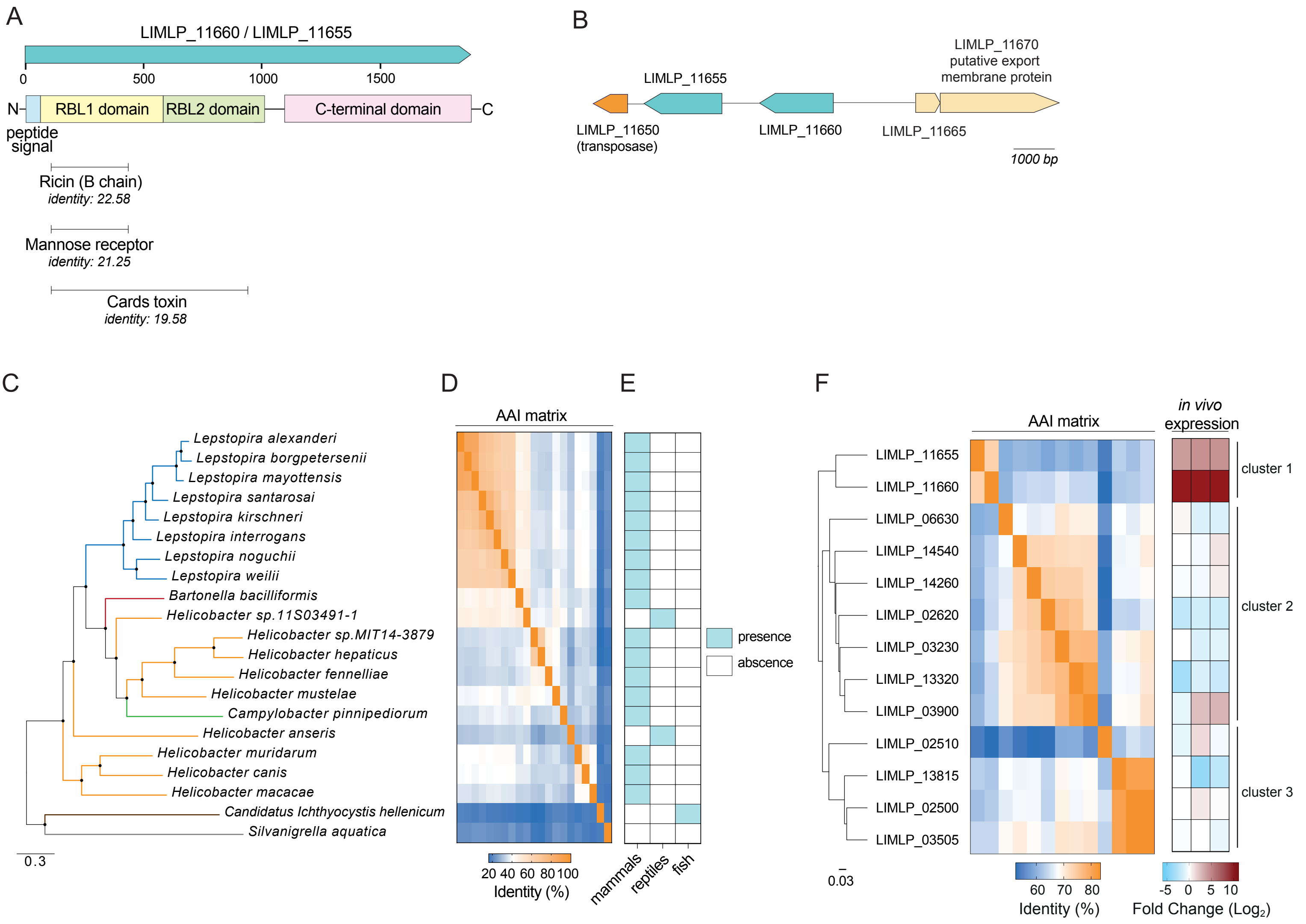


**Extended data Fig. 6**

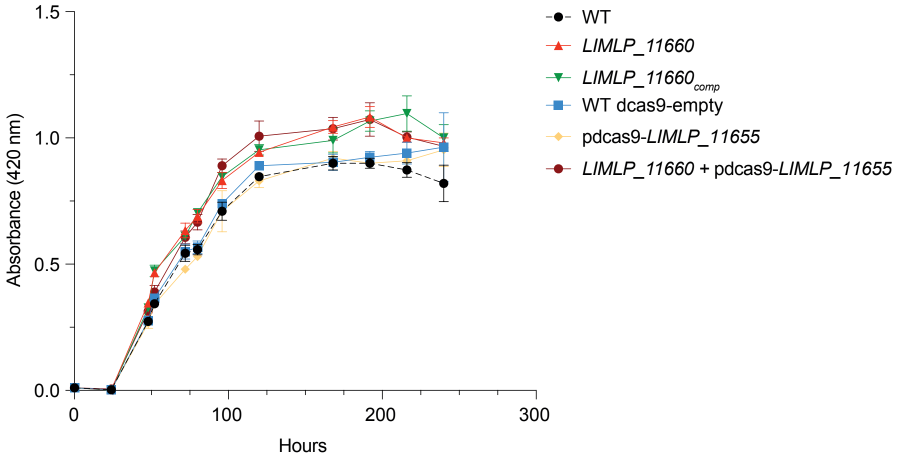


**Extended data Fig. 7**

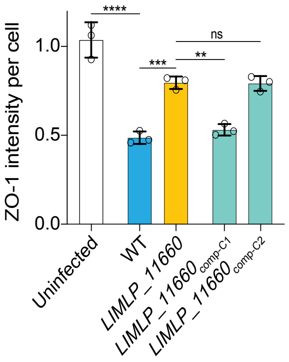


**Extended data Fig. 8**

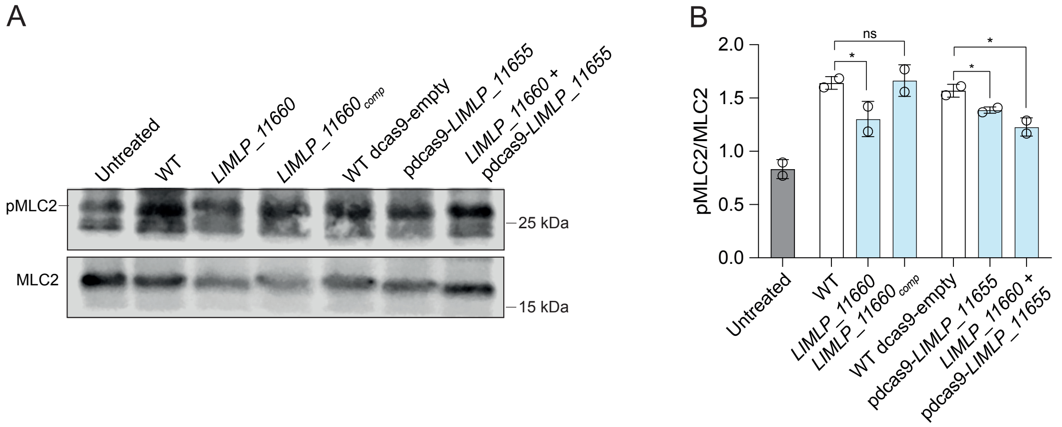


**Extended data Fig. 9**

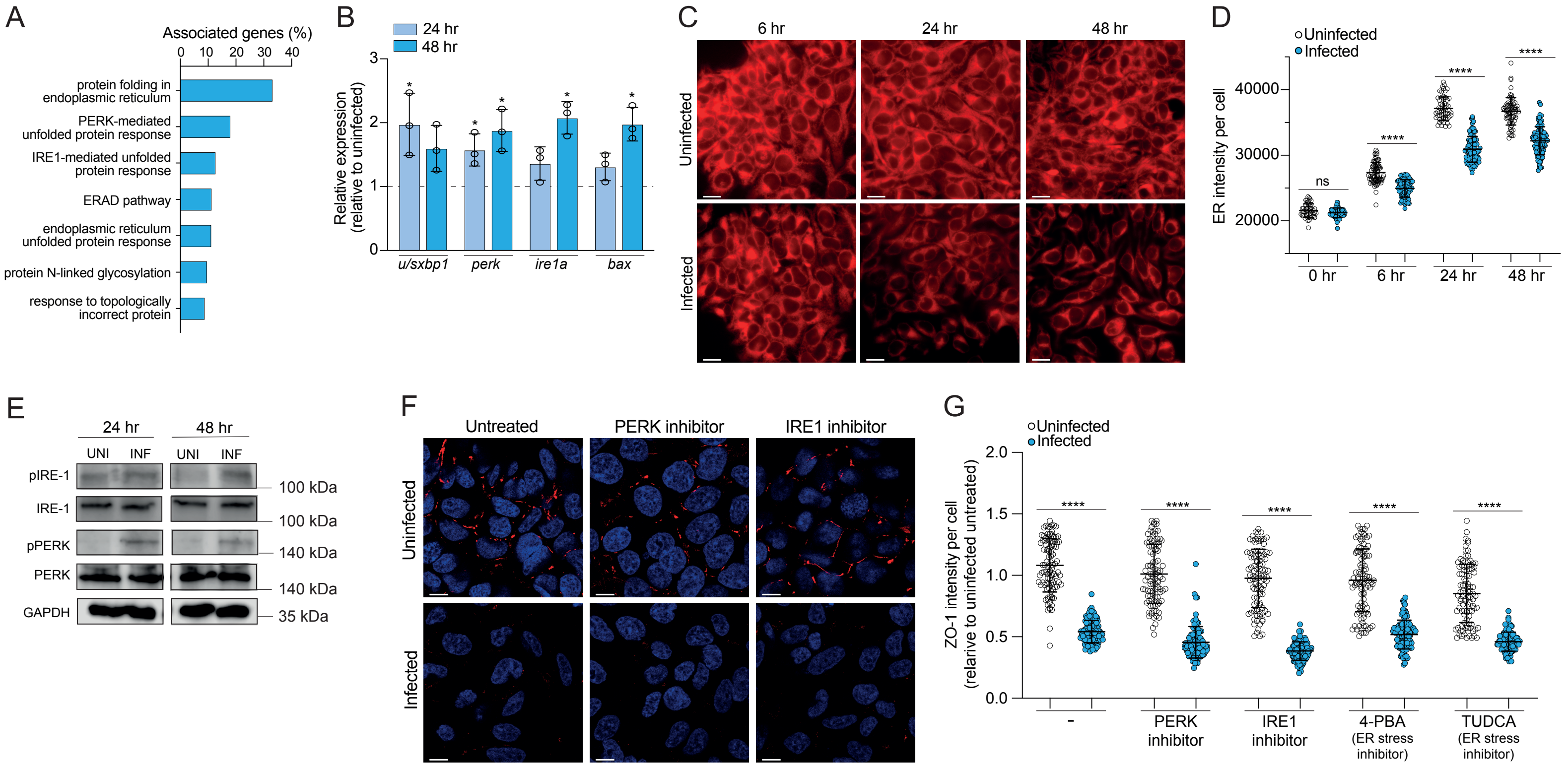


**Supplementary Fig. 1**

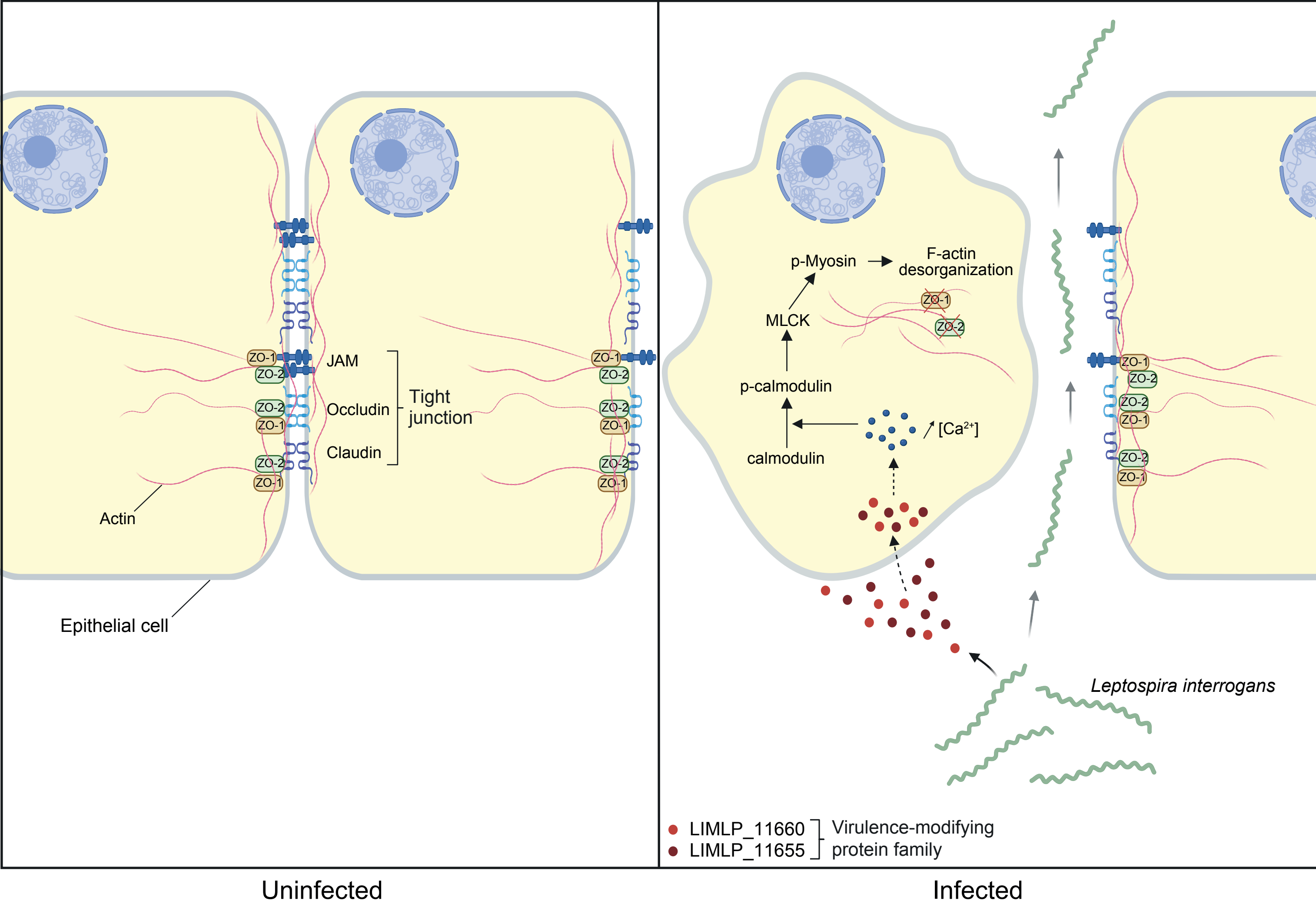


**Supplementary Fig. 2**
